# MegaLMM: Mega-scale linear mixed models for genomic predictions with thousands of traits

**DOI:** 10.1101/2020.05.26.116814

**Authors:** Daniel E Runcie, Jiayi Qu, Hao Cheng, Lorin Crawford

**Affiliations:** Department of Plant Sciences, University of California Davis, Davis, CA, USA; Department of Animal Sciences, University of California Davis, Davis, CA, USA; Microsoft Research New England, Cambridge, MA, USA

**Keywords:** Multi-trait Linear Mixed Model, Genomic prediction, High-throughput phenotyping, Multi-environment trial

## Abstract

Large-scale phenotype data can enhance the power of genomic prediction in plant and animal breeding, as well as human genetics. However, the statistical foundation of multi-trait genomic prediction is based on the multivariate linear mixed effect model, a tool notorious for its fragility when applied to more than a handful of traits. We present `MegaLMM`, a statistical framework and associated software package for mixed model analyses of a virtually unlimited number of traits. Using three examples with real plant data, we show that `MegaLMM` can leverage thousands of traits at once to significantly improve genetic value prediction accuracy.

## Background

New high-throughput phenotyping technologies hold promise for a revolution in data-driven decisions in plant and animal breeding programs (Araus *et al.* 2018; Koltes *et al.* 2019). For example, drone-based hyperspectral cameras can image fields at high resolution across hundreds of spectral bands (Rutkoski *et al.* 2016), wearable sensors can continuously monitor animals health and physiology (Neethirajan 2017), and RNA sequencing and metabolite profiling can simultaneously assay the concentrations of tens-of-thousands of targets (Schrag *et al.* 2018). These high-dimensional traits could allow breeders to rapidly assess many aspects of performance more accurately or earlier in development than was possible using traditional tools. This can increase the rate of gain in target traits by increasing selection accuracy, increasing selection intensity, and reducing breeding cycle durations.

However, efficiently incorporating high-dimensional phenotype data into breeding decisions is challenging. Whenever two traits are genetically correlated, joint analyses can improve the precision of variety evaluation (Thompson and Meyer 1986). However, two key problems emerge. First, the number of traits in high-dimensional datasets is often much larger than the number of breeding lines, which means that naive correlation estimates are not robust. Second, phenotypic correlation among traits are often poor approximations to genetic correlation, so not all correlated traits are useful for breeding decisions (Bernardo 2010). For example, plants grown in more productive areas of a field will tend to produce higher yields and be greener (measured by hyperspectral reflectance). Yet, selecting indirectly based on green plants instead of directly on higher yields may be counter-productive because “green-ess” may indicate an over-investment in vegetative tissues at the expense of seed. This contrasts with the problem of predicting genetic values from genotype data (e.g., genomic prediction; Meuwissen *et al.* (2001)), where all correlations between candidate features and performance are useful for selection.

The multivariate linear mixed model (MvLMM) is a widely-used statistical tool for decomposing phenotypic correlations into genetic and non-genetic components. The MvLMM is a multi-outcome generalization of the univariate linear mixed model (LMM) that forms the backbone of the majority of methods in quantitative genetics. The MvLMM was introduced over 40 years ago (Henderson and Quaas 1976), and has repeatedly been shown to increase selection efficiency (Piepho *et al.* 2007; Calus and Veerkamp 2011; Jia and Jannink 2012). Yet, MvLMMs are still rarely used in actual breeding programs because naive implementations of the framework are sensitive to noise, prone to overfitting, and exhibit convergence problems (Johnstone and Titterington 2009). Furthermore, existing algorithms are extremely computationally demanding. The fragility of naive MvLMMs is due to the number of variance-covariance parameters that must be estimated which increases quadratically with the number of traits. The computational demands increase even more dramatically: from cubically to quintically with the number of traits (Zhou and Stephens 2014) because most algorithms require repeated inversion of large covariance matrices. These matrix operations dominate the time required to fit a MvLMMs, leading to models that take days, weeks, or even years to converge.

Here, we describe `MegaLMM` (linear mixed models for millions of observations), a novel statistical method and computational algorithm for fitting massive-scale MvLMMs to large-scale phenotypic datasets. Although we focus on plant breeding applications for concreteness, our method can be broadly applied wherever multi-trait linear mixed models are used (e.g., human genetics, industrial experiments, psychology, linguistics, etc.). `MegaLMM` dramatically improves upon existing methods that fit low-rank MvLMMs, allowing multiple random effects and un-balanced study designs with large amounts of missing data. We achieve both scalability and statistical robustness by combining strong, but biologically motivated, Bayesian priors for statistical regularization–analogous to the *p >> n* approach of genomic prediction methods–with algorithmic innovations recently developed for LMMs. In the three examples below, we demonstrate that our algorithm maintains high predictive accuracy for tens-of-thousands of traits, and dramatically improves the prediction of genetic values over existing methods when applied to data from real breeding programs.

## Results

### Methods overview

`MegaLMM` fits a full multi-trait linear mixed model (MvLMM) to a matrix of phenotypic observations for *n* genotypes and *t* traits (level 1 of Figure 1A). We decompose this matrix into fixed, random, and residual components, while modeling the sources of variation and covariation among all pairs of traits. The main statistical and computational challenge of fitting large MvLMMs centers around the need to robustly estimate *t × t* covariance matrices for the residuals and each random effect. Each covariance matrix has *t*(*t* − 1)/2 + *t* free parameters, and any direct estimation approach is computationally demanding because it requires repeatedly inverting these matrices (an 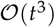 operation).

**Figure 1.**
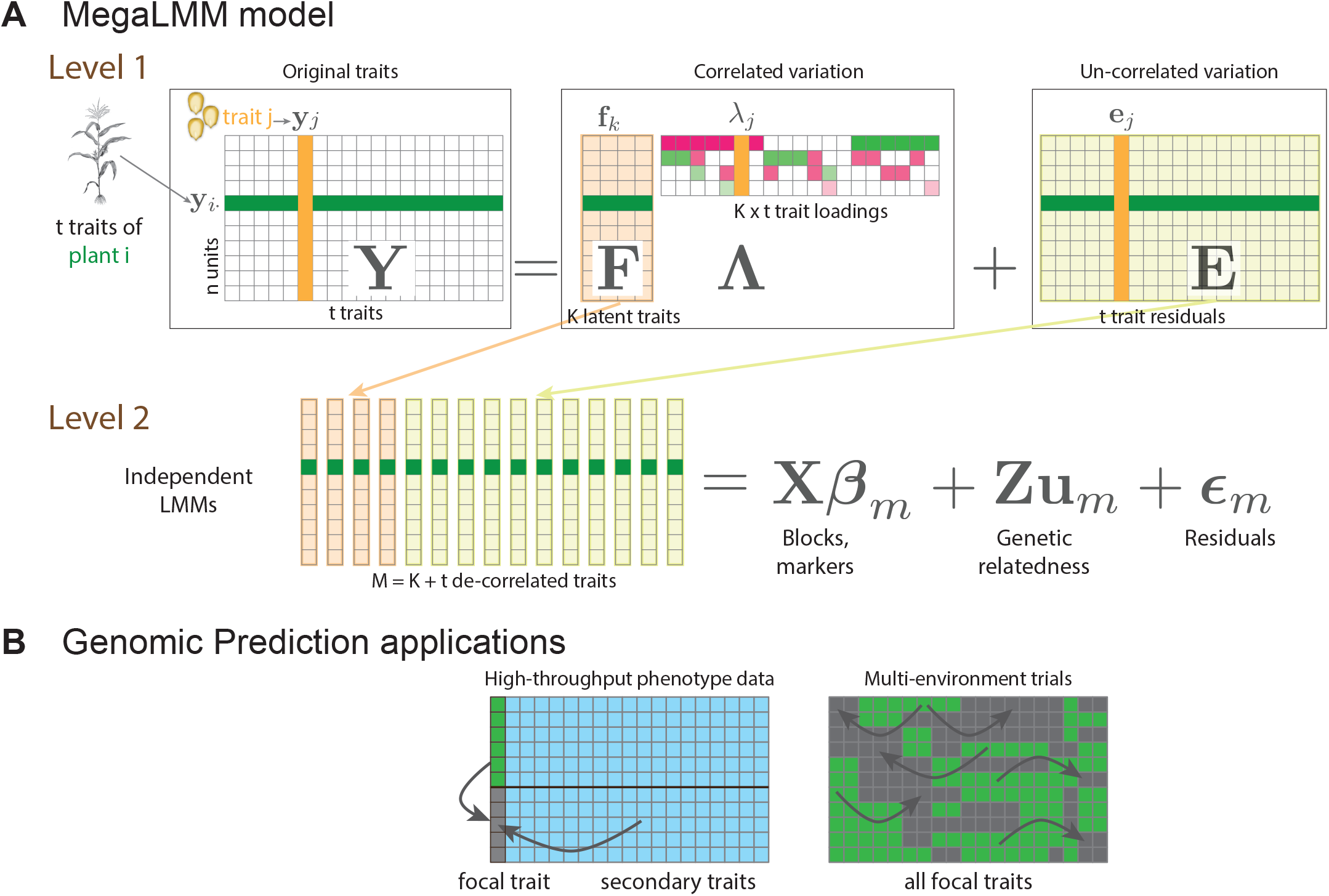
Overview of the MegaLMM model: **A.**`MegaLMM` decomposes a typical MvLMM into a two-level hierarchical model. In level 1, raw data from *t* traits on each of the *n* plants (more generally observational units) (**y**_*i*_) are combined into an *n* × *t* trait matrix **Y**. Variation in **Y** is decomposed into two parts: a low-rank model (**FΛ**) consisting of *K* latent factor traits, each of which controls variation in a subset of the original traits through the loadings matrix **Λ**, and a residual matrix (**E**) of independent residuals for each trait. The latent factor traits and the *t* residual vectors are now mutually un-correlated, and are each modeled with independent LMMs in level 2. Experimental design factors, genetic background effects, and other modeling terms are introduced at this level. Cells highlighted in green show observations and associated parameters for plant *i*. Cells highlighted in orange highlight observations and associated parameters for trait *j*. **B.** Two multi-trait genomic prediction applications: i) the use of high-throughput phenotyping data to supplement for expensive direct measures of focal traits like grain yield, and ii) the analysis of large multi-environment trials. In each case, observed data of focal traits (green) and secondary traits (blue) are used to predict genetic values for individuals without direct phenotypic observations (grey).

We solve both of these problems by introducing *K* un-observed (latent) traits called factors (**f**_*k*_) to represent the causes of covariance among the *t* observed traits. We treat each latent trait just as we would any directly measured trait and decompose its variation into the same fixed, random and residual components using a set of parallel univariate linear mixed models (level 2 of Figure 1A). We then model the pairwise correlations between each latent trait and each observed trait through *K* loadings vectors ***λ**_k_*.

Together, the set of parallel univariate LMMs and the set of factor loading vectors result in a novel and very general re-parameterization of the MvLMM framework as a mixed-effect factor model. This parameterization leads to dramatic computational performance gains by avoiding all large matrix inversions. It also serves as a scaffold for eliciting Bayesian priors that are intuitive and provide powerful regularization which is necessary for robust performance with limited data. Our default prior distributions encourage: i) shrinkage on the factor-trait correlations (*λ_jk_*) to avoid over-fitting covariances, and ii) shrinkage on the factor sizes to avoid including too many latent traits. This two-dimensional regularization helps the model focus only on the strongest, most relevant signals in the data.

While others have used latent factor approaches to reduce dimensionality of MvLMMs (e.g., de Los Campos and Gianola 2007; Meyer 2007; Runcie and Mukherjee 2013; Dahl *et al.* 2016), these methods only use factors for a single random effect (usually the matrix of random genetic values)–with the exception of BSFG which uses factors for the combined effect of a single random effect and the residuals (Runcie and Mukherjee 2013). In `MegaLMM`, we expand this framework and use factors to model the joint effects of all predictors: fixed, random and residual factors on multiple traits.

We combine this efficient factor model structure with algorithmic innovations that greatly enhance computational efficiency, drawing upon recent work in LMMs (Kang *et al.* 2008; Zhou and Stephens 2012; Lippert *et al.* 2011; Runcie and Crawford 2019). While Gibbs samplers for MvLMMs are notoriously slow, we discovered extensive opportunities for collapsing sampling steps, marginalizing over missing data, and discritizing variance components so that intermediate results can be cached (Supplemental Methods).

Genomic prediction using `MegaLMM` works by fitting the model to a partially observed trait matrix, with the traits to be predicted imputed as missing data. `MegaLMM` then estimates genetic values for all traits (both observed and missing) in a single step (Figure 1B).

### MegaLMM is efficient and effective for large datasets

We used a gene expression matrix with 20,843 genes measured in each of 665 *Arabidopsis thaliana* accessions (a total of nearly 14 million observations), to evaluate the accuracy and time requirements for trait-assisted genomic prediction–a classic example of an applied use of MvLMMs–across a panel of existing software packages. We created datasets with 4 to 20,842 “secondary” traits with complete data, and used these data to predict the genetic values of a single randomly selected “focal” gene with 50% missing data.

Despite the limited number of independent lines in this data set, adding up to ≈200 secondary traits improved the genomic prediction accuracy of `MegaLMM` and two other Bayesian methods: `MCMCglmm` and phenix (Figure 2A). The maximum likelihood method `MTG2` (Lee and van der Werf 2016), on the other hand, did only marginally better than single-trait prediction, and genomic prediction accuracy declined with 32 traits, likely due to overfitting. We note that the results here are averages over 20 randomly selected focal genes. The prediction accuracy and benefits of multi-trait prediction varied considerably among genes (Figures S1 and S2), but comparisons among methods were largely correlated. Using simulated datasets where we knew the true genetic and residual covariances among traits, we also found that `MegaLMM` was at least as accurate in estimating covariance parameters as the competing methods (Figure S3).

**Figure 2.**
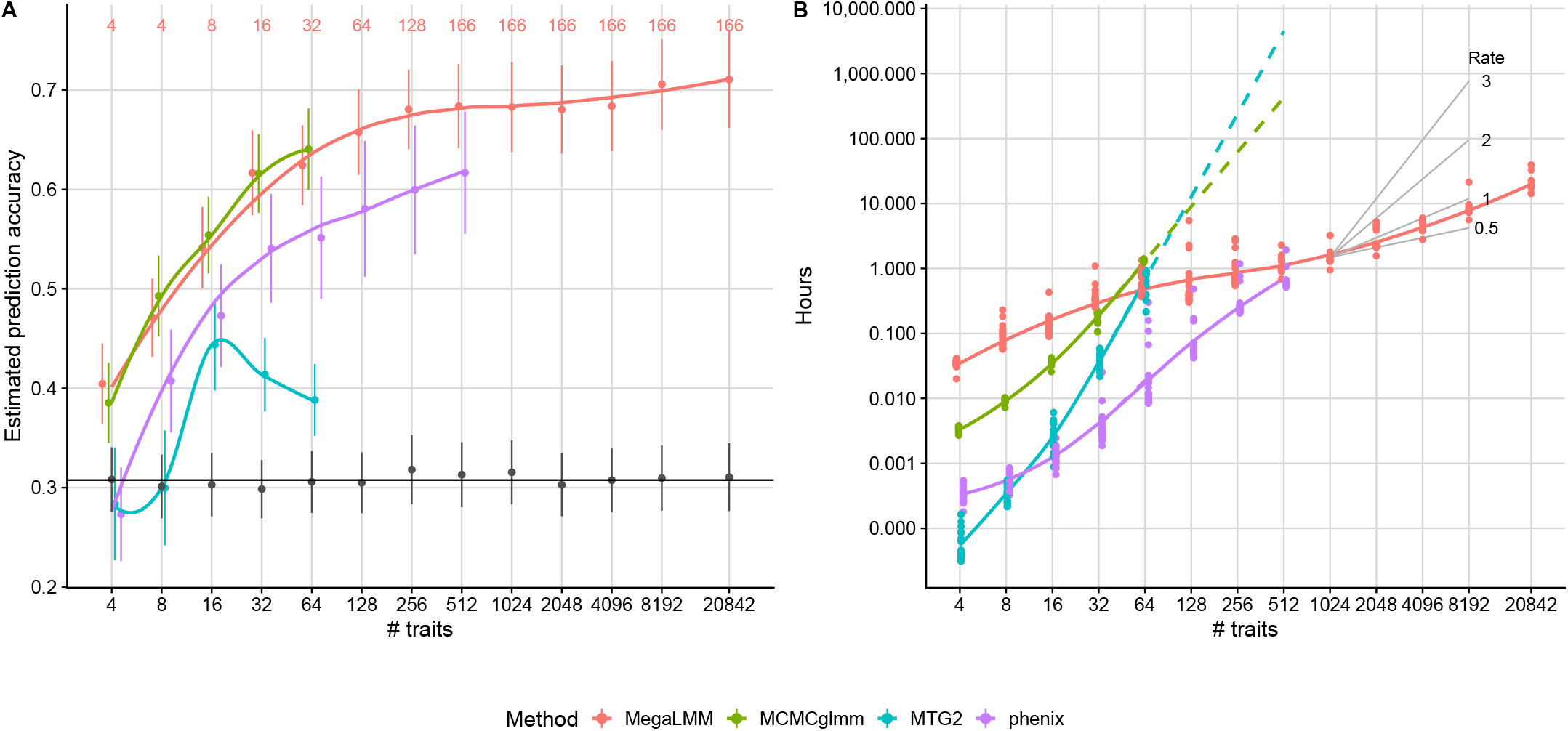
MegaLMM scales efficiently for very high-dimensional traits. Four competing methods were used to fit multi-trait genomic prediction models to predict genetic values for a single focal gene expression trait using complete data from *t* additional traits. Data are from an *Arabidopsis thaliana* gene expression data with 20,843 genes and 665 lines. **A)** Average estimated genomic prediction accuracy across 20 focal traits using *t* additional secondary traits for each of the four prediction methods (the horizontal line is the average univariate prediction accuracy). Genomic prediction accuracy was estimated by cross-validation as 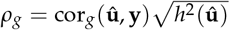 to account for non-genetic correlations between the secondary traits and focal traits since all were measured in the same sample. Smoothed curves are estimated by stats::lowess. The number of latent factors used for `MegaLMM` (*K*) is listed in red at the top of the figure. **B)** Computational times required to find a solution for each MvLMM. For the MCMC methods `MCMCglmm` and `MegaLMM`, times were estimated as the time required to collect an effective sample size of at least 1000 for *>* 90% of the elements in the genetic covariance matrix **U**. Computational times for `MCMCglmm` and MTG2 above 64 traits were linearly extrapolated (on log scale) based on the slope between 32 and 64 traits. Black lines show the slope of exponential scaling functions with the specified exponents for reference.

Beyond 32 secondary traits, computational times for `MCMCglmm` and `MTG2` became prohibitive (Figure 2B). Using extrapolation, we estimated that fitting these methods for 512 traits would take 20 days and 217 days, respectively, without considering issues of model convergence. In contrast, `phenix` and `MegaLMM` were both able to converge on good model fits for 512 traits in approximately one hour.

Beyond 512 traits, `MegaLMM` was the only viable method as phenix cannot be applied to datasets with *t > n* phenotypes. Although the genomic prediction accuracy of `MegaLMM` did not increase further after ≈256 traits, performance did not suffer even with the full dataset of *>* 20, 000 traits and the analysis was completed in less than a day. This shows that `MegaLMM` is feasible to apply to very high-dimensional studies and, in most cases, does not require pre-filtering of traits–something that requires great care in genomic prediction applications to avoid misleading results (Runcie and Cheng 2019).

An important feature of `MegaLMM` is that the choice of the number of latent factors *K* is less critical than in most factor models. Since factors are ordered from most-to-least important by the prior (See Methods), as long as enough factors are specified to capture the majority of the covariance among traits, adding additional latent factors does not lead to over-fitting (Figure S4A). Additional factors do increase the run-time of the algorithm, though (Figure S4B), so some optimization of *K* during the burn-in period can reduce computational demands during posterior sampling.

### Applications to real breeding programs

To demonstrate the utility of `MegaLMM`, we developed two classes of genomic prediction models for high-dimensional phenotype data in real plant breeding programs.

### Genomic prediction using hyperspectral reflectance data

When the final performance of a variety is difficult or costly to obtain, breeding programs can supplement direct measures of performance with data from surrogate traits that can be measured cheaply, earlier in the breeding cycle, and on more varieties. For example, in the bread wheat breeding program at CIM-MYT, hyperspectral reflectance data can be collected rapidly and repeatedly by aerial drones on thousands of plots (Krause *et al.* 2019). We developed a multi-trait genomic prediction model to incorporate 62-band hyperspectral reflectance data from 10 different drone flights over the course of one growing season, and compared the accuracy of these genetic value predictions against more traditional approaches.

We first compared three standard univariate methods: `GBLUP` (Hayes *et al.* 2009), `Bayesian LASSO (BL)` (Park and Casella 2013), and Reproducing kernel Hilbert space (`RKHS`) regression (de Los Campos *et al.* 2010). `GBLUP` achieved a prediction accuracy of *ρ_g_* = 0.43 for yield (Figure 3A). Both the `BL` and `RKHS` methods showed modest improvements, with *ρ_g_* = 0.47 and *ρ_g_* = 0.49, respectively in these data. The `RKHS` model often out-performs `GBLUP` in plant breeding datasets, but improvements are generally slight and inconsistent depending on the genetic architecture of the targeted trait.

**Figure 3.**
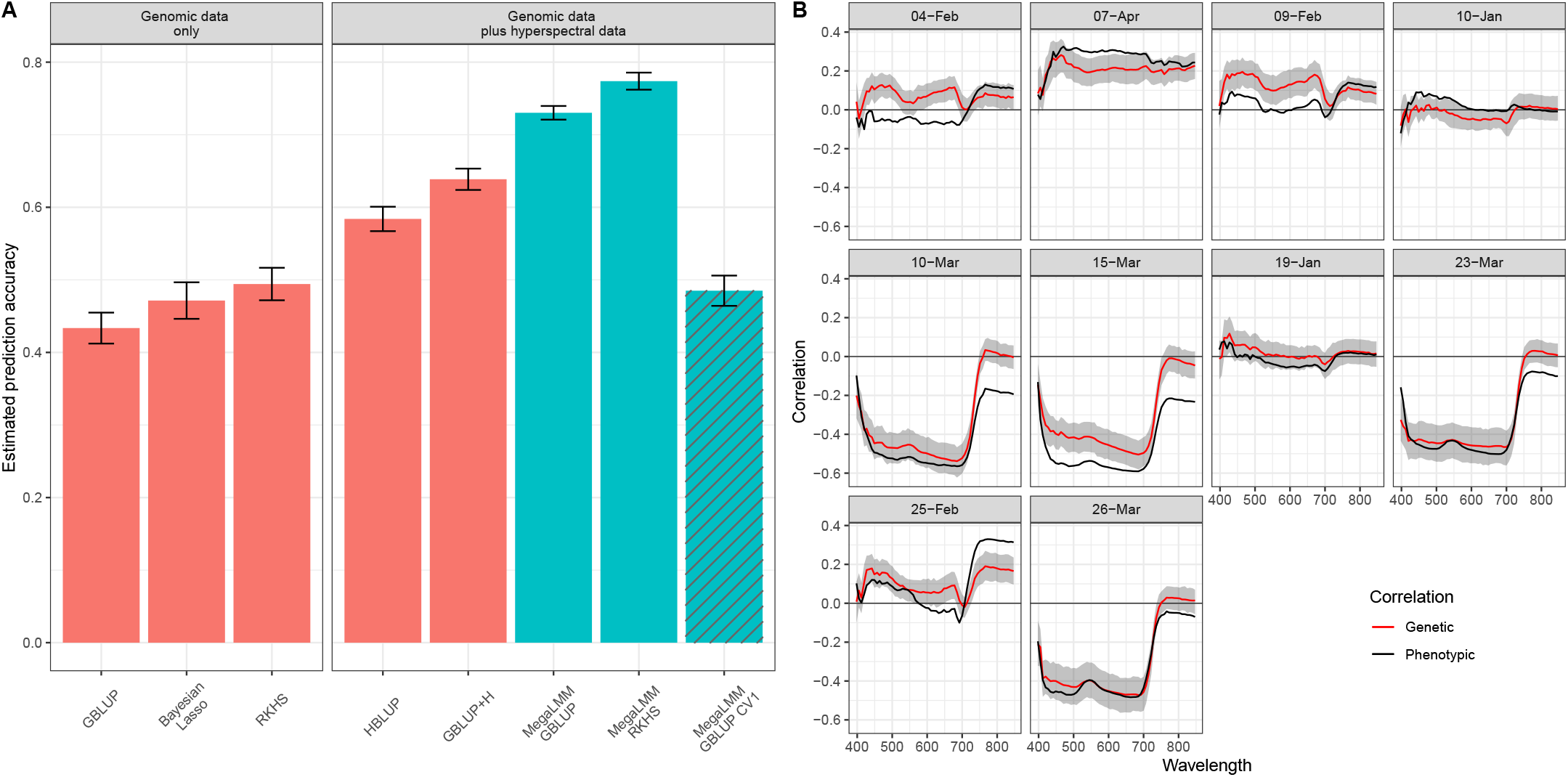
Performance of single-trait and multi-trait genomic prediction for wheat yield. **A)**8 methods for predicting Grain Yields of 1,092 bread wheat lines. Genetic value prediction accuracy was estimated by cross-validation. Complete data (yield, marker genotypes, and 620 hyperspectral wavelength reflectances) was available for all lines, but 50% of the yield values were masked during model training. Genetic value prediction accuracy was estimated as 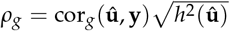 because hyperspectral data and actual yields were collected on the same plots (Runcie and Cheng 2019). Bars show average estimates (± standard error) over 20 replicate cross-validation runs for each method. Details of each model are presented in the Supplemental Methods. Briefly, the three single-trait methods only used yield and genotype data. The five multi-trait methods additionally used hyperspectral data measured on all 1,092 lines. **B)** Phenotypic correlation (black lines), and estimates of genetic correlation (red lines) between each hyperspectral wavelength measured on each of the 10 flight dates with final grain yield. Genetic correlations were estimated with the `MegaLMM` GBLUP method using complete data. Ribbons show the 95% highest posterior density (HPD) intervals.

In the original analysis of this dataset, Krause *et al.* (2019) achieved increased performance by replacing the genomic kernel (**K** in our notation) with a kernel based on the cross-product of hyperspectral reflectances across all wavelengths and time points (termed the **H** matrix). We replicated these results, achieving a prediction accuracy of *ρ_g_* = 0.58 (`HBLUP` method). These authors also proposed a multi-kernel model combining the **K** and **H** kernel matrices, although they only applied this to cross-treatment genotype-by-environment predictions. We found that applying this multi-kernel method to the within-environment data resulted in additional accuracy gains (*ρ_g_* = 0.64) (`GBLUP+H` method; Figure 3A).

While more effective than univariate methods, predictions based on the **H** kernel matrix are biased by non-genetic correlations between surrogate traits and yield because they do not directly model the genetic component of these correlations. `MegaLMM` implements a full multi-trait mixed model and thus can separate these sources of correlation. We fit three different multi-trait prediction models with `MegaLMM`. The first was a standard multi-trait mixed model with a single random effect based on the genomic relationship matrix **K**. This method achieved a dramatically higher prediction accuracy than any of the previous approaches (*ρ_g_* = 0.73). Second, because the RKHS model had the highest performance among univariate predictions, we implemented an approximate RKHS method in `MegaLMM` based on averaging over three kernel matrices (de Los Campos *et al.* 2010). We are not aware of any other high-dimensional MvLMM implementations that allow models with multiple random effects. This model achieved the highest predictive accuracy (*ρ_g_* = 0.77). Finally, we repeated the `MegaLMM-GBLUP` analysis but this time masking all phenotype data (both grain yield and hyperspectral data) from the testing set. We called this approach `MegaLMM-GBLUP-CV1` following the nomenclature from Burgueño *et al.* (2012). Genetic prediction accuracy in the CV1 setting was similar to the univariate methods (*ρ_g_* = 0.49), showing that nearly all benefit of `MegaLMM` in this dataset came through the optimal use of secondary trait phenotypes on the lines of interest.

In summary, by directly modeling the genetic covariance between the surrogate traits (hyperspectral reflectance measures), we achieved performance increases of 56%-79%, and up to 36% over the HBLUP method. To show that these conclusions were robust in other datasets, we repeated the same analyses in the other 19 trials reported by Krause *et al.* (2019) and results were highly similar in all trials (Figure S5).

To explore *why* directly modeling the genetic correlation is important, we compared the estimated genetic correlations between each hyperspectral band and grain yield to the corresponding phenotypic correlations (Figure 3B). Most genetic correlation estimates closely paralleled the phenotypic correlations, with the largest values for low-to-intermediate wavelengths occurring on dates towards the end of the growing season while plants were in the grain filling stage (Krause *et al.* 2019). However, there were notable differences. For example, genomic correlations were moderate (*ρ_g_* ≈ 0.2) for most wavelengths during early February sampling dates while phenotypic correlations were near zero; yet, during early March time points, phenotypic correlations between yield and bands around 800 nanometers were moderate (*ρy* ≈ − 0.2) but genomic correlations were near-zero. `MegaLMM` is able to model the discrepancy between genomic and phenotypic correlations, but methods based on the **H** matrix (e.g., `HBLUP`) are not.

### Genomic prediction of agronomic traits across multi-environmental trials

Multi-trait mixed models are also used to analyze data from multi-environment trials to account for genotype-environment interactions and select the best genotypes in each environment. The Genomes2Field initiative (https://www.genomes2fields.org/) is an ongoing multi-environment field experiment of maize hybrid genotypes across 20 American states and Canadian provinces. Data from the years 2014-2017 included 119 trials with a total of 2102 hybrids. As in many large-scale multi-environment trials, only a small proportion of the available genotypes were grown in each trial. Therefore, the majority of trial-genotype combinations were un-observed.

We selected four representative agronomically important traits and compared the ability of four modeling approaches to impute the missing measurements. Including across-trial information was beneficial for each of the four traits, suggesting generally positive genetic correlations across trials. However, applying `MegaLMM` to each of the four trait datasets improved predictions dramatically, with average benefits across trials ranging from *ρ_y_* = 0.10 to *ρ_y_* = 0.17 (Figure 4). The performance of `phenix` was inconsistent across traits and trials, likely because its model for the non-additive genetic covariance (i.e., the residual) is less flexible than `MegaLMM`.

**Figure 4.**
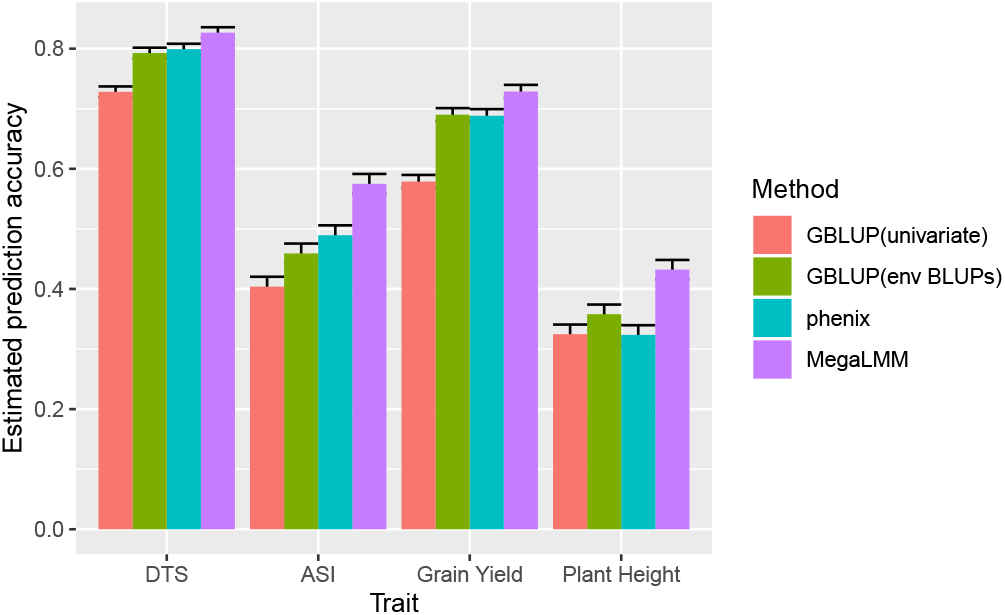
Average within-trial prediction accuracy for four maize traits in the Genomes2Fields Initiative experiment. Traits included: days to silking (DTS), anthesis-silking interval (ASI), grain yield, and plant height. Bars show the average ± 95% confidence intervals of prediction accuracy for each method across the 76-99 trials with sufficient training data for each trait. For each trail, prediction accuracies were estimated as the mean over 20 randomized cross-validation replicates.

To explore *why* jointly modeling all genetic and non-genetic covariances for each pair of trials improved prediction accuracy for each trait, we assessed the per-trial differences in performance between `MegaLMM` and the corresponding within-trial genomic prediction model. Trials varied considerably in how much `MegaLMM` improved genomic prediction accuracy, with several trials seeing improvements of *ρ >* 0.4. The magnitude of the `MegaLMM` effect on genomic prediction accuracy was largely explained by the maximum genetic covariance between that trial and any other trial in the dataset (Figure S6). This is expected because the benefit of a MvLMM is largely dependent on the magnitude of genetic covariances between traits.

A common approach in multi-environment trials is to combine similar trials (based on geographic location or similar environments) into clusters and make genetic value predictions separately for each cluster (Piepho and Möhring 2005). However, this will not be successful if clusters cannot be selected *a priori* because using the trial data itself to identify clusters can lead to overfitting if not performed carefully (Runcie and Cheng 2019). In these data, the distribution of genomic correlations between trials differed among traits, so it is not straightforward to identify which pairs or subsets of trials could be combined. The most obvious predictor of trial similarity is geographic distance, but we did not see consistent spatial patterns in the among-trial covariances across the four traits. The trials with the greatest benefit from our MvLMM showed geographic clustering in the central mid-west for the anthesis-silking interval (ASI) but not for the other three traits (Figure 5A). Genetic correlations tended to decrease over long distances for ASI and over short distances for plant height, but not for the other two traits (Figure 5B), resulting in obvious geographic clustering of genetic correlations for ASI but not the other traits (Figure 5C). This suggests that including all trials together in one model is necessary to maximize the benefit of the MvLMM approach to multi-environment plant breeding.

**Figure 5.**
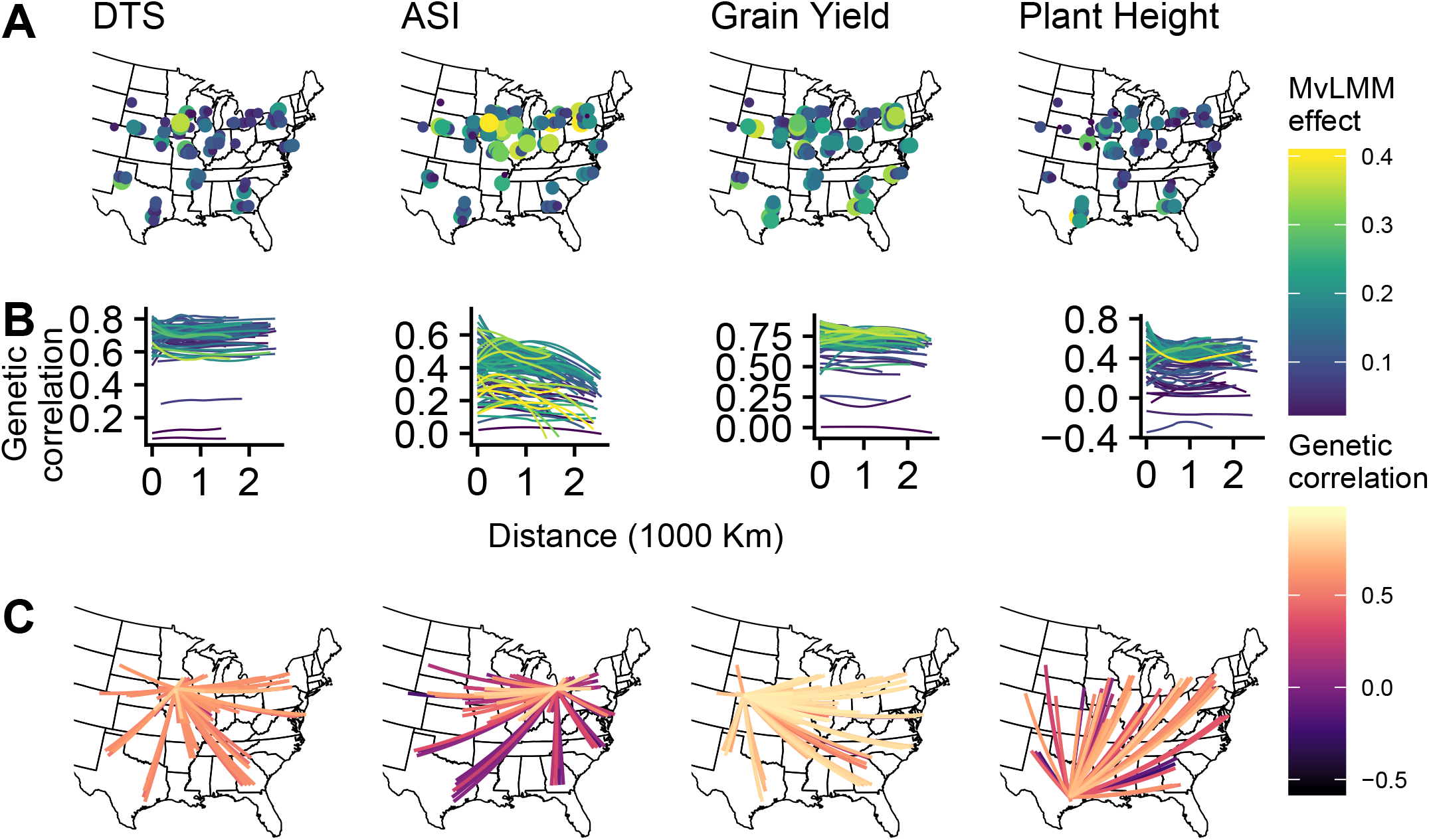
Benefit of *MegaLMM* and geographic distributions of among-trial genetic correlations vary among traits. Traits analyzed included: days to silking (DTS), anthesis-silking interval (ASI), grain yield, and plant height. **A**) Trial locations for each trait are shown. Points were jittered west-to-east to prevent overlap of repeated trials across years. Size and color of each point correspond to the increase in prediction accuracy for `MegaLMM` versus a univariate LMM. **B**) Smoothed estimates (computed using `geom_smooth` with a bandwidth of 1.0) of the relationship between geographic distance and genetic correlation for each trial. Line colors correspond to the benefit of `MegaLMM` in each focal trial. **C**) Genetic correlations between the trial with the greatest benefit of `MegaLMM` for each trait and each other trial.

## Discussion

Novel statistical methods can help optimize plant and animal breeding programs to meet future food security needs. In the above examples, we highlighted two areas where large-scale phenotype data can improve the accuracy of genomic prediction in realistic plant breeding scenarios: by incorporating high-throughput phenotyping data from remote sensors, and by synthesizing data on gene-environment interactions across large-scale multi-environment trials. In both examples, we apply high-dimensional multivariate linear mixed models to efficiently integrate all available genotype and phenotype data into genetic value predictions. `MegaLMM` is a scalable tool that extends the feasible range of input data for multivariate linear mixed models by at least two orders of magnitude over existing methods, while providing the flexibility to plug directly into existing breeding programs.

### Computational and statistical efficiency

Computational issues in single-trait LMMs have been studied extensively, allowing implementations for large datasets (Lip-pert *et al.* 2011; Zhou and Stephens 2014; Loh *et al.* 2015; Runcie and Crawford 2019). Most of these algorithms diagonalize the genomic relationship matrices to improve computational efficiency. This technique dramatically improves the performance of simple, low-dimensional MvLMMs as well (e.g., Zhou and Stephens 2014; Lee and van der Werf 2016). However, diagonalization does not address the computational challenge imposed by large trait-covariance matrices, and can only be applied to models with a single random effect and no missing data. Therefore, these tools cannot be applied to the datasets studied here or, more generally, to most large-scale studies of gene-environment interactions that frequently have large proportions of missing data (Piepho *et al.* 2007) (Figure 1) and to studies that have experimental designs with multiple sources of covariance (e.g., spatial environmental variation or non-additive genetics).

Our work builds on the factor-analytic approach to regularizing MvLMMs (de Los Campos and Gianola 2007; Meyer 2007; Runcie and Mukherjee 2013; Dahl *et al.* 2016) and is most similar to `BSFG` (Runcie and Mukherjee 2013) and `phenix` (*Dahl et al.* 2016), which improve upon traditional quantitative genetic factor models by specifying sparse or low-rank factor matrices to improve robustness in high dimensions. Importantly, however, these models are limited to a single random effect and are not tractable for datasets with large numbers of traits because of computational inefficiencies (`BSFG`), or a lack of strong regularization on the residual covariance matrix (`phenix`). `MegaLMM` generalizes both methods and dramatically improves their weaknesses, allowing analyses with >20,000 traits to be completed in less than one day. Since `MegaLMM` scales approximately linearly with the number of traits (Figure 2), applying it to datasets with many more traits may be feasible. While we have designed many of our routines to take advantage of multi-core CPUs, graphical processing units may offer additional performance gains.

Two key advantages of `MegaLMM` are its flexibility and generality. We have designed the `MegaLMM R` package to be as general as possible so that it can be applied to a wide array of problems in quantitative genetics. `MegaLMM` tolerates unbalanced designs with incomplete observations (something that makes `MCMCglmm` and `MTG2` very slow), arbitrarily complex fixed effect specifications to model experimental blocks, covariates, or other sources of variation among samples (unlike `phenix`), and most importantly, multiple random effects (unlike `phenix`, `GEMMA`, or `MTG2`). Multiple random effect terms can be used to account for spatially correlated variation across fields, non-additive genetic variation that is not useful for breeding, or to more flexibly model non-linear genetic architectures as we demonstrated with the approximate RKHS regression approach in the wheat application (Figure 3). To make multiple-random-effect models computationally efficient, we take our earlier work with LMMs (Runcie and Crawford 2019) and extend the same discrete estimation procedure to MvLMMs where the impact on computational efficiency is exponentially greater. Other commonly used tools for fitting MvLMMs such as ASREML (Gilmour 2007) allow more flexibility in the specification of multiple variance-component models with correlated random effects that are not currently possible in `MegaLMM`. However, these tools do not scale well beyond ≈10 traits, so are not feasible to apply directly to large-scale datasets in plant breeding.

### Applicability to high-throughput phenotypic data

Large-scale phenotype data collection is rapidly emerging as a standard tool in plant breeding and other fields that use quantitative genetics (GTEx Consortium 2017; Araus *et al.* 2018; Bycroft *et al.* 2018). These deep phenotyping datasets can be used as high-dimenisional features to predict genetic values in agronomically important traits and serve as substitutes for direct assays where these are more time-consuming or expensive to collect.

Breeding objectives differ from the goals of polygenic risk score predictions for human diseases because the prediction target is not the phenotype of an individual, but its genetic value (Runcie and Cheng 2019). Genetic values quantify the expected phenotype of a plant’s offspring, and so exclude impacts of the plant’s own microenvironment on its phenotype (Bernardo 2010). Therefore, accurate genetic value prediction requires models that can distinguish between genetic and non-genetic sources of covariation among traits.

The MvLMM is considered the gold-standard method for isolating genetic correlations from non-genetic correlations in genetic value prediction (Piepho *et al.* 2007). However, it has rarely been applied in breeding programs because of the computational challenges associated with estimating multiple large covariance matrices. With high-throughput phenotype (HTP) data, MvLMMs have only been applied directly to sets of ≈2 − 5 traits. Instead, several authors have used a prior round of feature selection or calculated summary statistics of the HTP to generate model inputs rather than using the raw high-dimensional data itself (e.g., Jia and Jannink 2012; Guo *et al.* 2014; Rutkoski *et al.* 2016; Sun *et al.* 2017; Crain *et al.* 2018). Other authors have replaced the MvLMM with a direct regression on the HTP data, using techniques such as factorial regression (van Eeuwijk *et al.* 2019), functional regression (Montesinos-López *et al.* 2017), kernel regression (Krause *et al.* 2019), and deep learning(Cuevas *et al.* 2019). While straightforward to implement, this conditioning on the HTP traits creates a form of collider bias which can induce genotype-phenotype associations that do not actually exist and impede genetic value predictions (Runcie and Cheng 2019). Alternative methods including IBCF (Juliana *et al.* 2019)) and regularized selection indexes (Lopez-Cruz *et al.* 2020) avoid computational complexities of the full MvLMMs, but do not make full use of the trait correlations in the data.

`MegaLMM`, on the other hand, fits a full MvLMM to an arbitrary number of HTP traits and should be more efficient at leveraging high-dimensional genetic correlations while accounting for non-genetic sources of covariance, particularly for datasets when HTP traits and focal performance traits are measured on the same plants. Non-genetic correlations will be less important on datasets where these sets of traits are measured on different plots. At least in the wheat breeding trial datasets we examined, the benefit of multi-trait modeling was much greater when traits were partially observed on each individual than when secondary traits were only observed in the training partition. This is expected theoretically and has been demonstrated previously in simulations Runcie and Cheng (2019), but the magnitude of the benefit was particularly dramatic here. This suggests that breeding programs should focus on developing HTP technologies that can measure secondary traits on the target individuals; HTP measurements on training individuals may be less useful for prediction applications. Unlike other methods, including too many traits, or including redundant traits that are highly correlated is unlikely to significantly impact prediction accuracy, reducing the need to carefully choose which traits to include and which to exclude *a priori*; `MegaLMM` allows users to simply include all traits they have at once.

### Applicability to multi-environment trial data

The analysis of multi-environment trials provides a separate set of computational and statistical challenges for plant breeders. Multi-environment trials (METs) are necessary because gene-environment interactions (GEIs) often prevent the same variety from performing best in all locations where a crop is grown (Piepho *et al.* 2007). However, METs are expensive and logistically difficult. Genomic predictions in METs could reduce the need to test every variety in every environment, allowing smaller individual trials (Heffner *et al.* 2009).

GEIs can be modeled in two ways: (i) as changes in variety effects on the same trait across environments (i.e., variety-by-environment interactions), or (ii) as a set of genetically correlated traits, with each trait-environment combination considered as a different phenotype (Piepho *et al.* 2007). When formulated with linear mixed models and random genetic effects, these two approaches are mathematically equivalent. Traditionally, the most common model for analyzing METs has been the AMMI model in which the genetic effects of each variety in each environment are modeled using a set of products between genetic and environmental vectors (Gauch 1988). AMMI models are used to rank genotypes in different environments and to identify environmental clusters with similar rankings of varieties. However, AMMI models cannot easily incorporate marker data. When genetic values are treated as random effects, AMMI models becomes factor models (generally called factor analytic models in this literature) (e.g. Piepho 1998; Smith *et al.* 2001), and can incorporate genetic marker data (e.g. Jarquín *et al.* 2014). `MegaLMM` extends this factor-analytic method for analyzing METs, making the methods robust for METs with hundreds or more individual trials.

A limitation of the AMMI factor-analytic approach to analyzing METs is that there is no mechanism for extending predictions to new environments outside of those already tested. Even large-scale commercial trials cannot test every field a farmer might use. Several authors have proposed using environmental covariates (ECs) to model environmental similarity in METs and predict GEIs for novel environments (e.g., Jarquín *et al.* 2014; Malosetti *et al.* 2016; Rincent *et al.* 2019). These approaches all involve regressions of genetic variation on the ECs, and so, if relevant ECs are missing or the relationship between variety plasticity and ECs is non-linear, these models will under-fit the GEIs. Nevertheless, these approaches are promising and have been successfully applied to large METs (e.g. Jarquín *et al.* 2014). `MegaLMM` cannot currently incorporate ECs to predict novel environments. However, a possible extension could involve replacing the *iid* prior on the elements of the factor loadings matrix with a regression on the ECs. This hybrid of ECs and a full MvLMM could leverage the strengths of both approaches.

### Model limitations

While `MegaLMM` works well across a wide range of applications in breeding programs, our approach does have some limitations.

First, since `MegaLMM` is built on the `Grid-LMM` framework for efficient likelihood calculations (Runcie and Crawford 2019), it does not scale well to large numbers of observations (in contrast to large numbers of traits), or large numbers of random effects. As the number of observational units increases, `MegaLMM`’s memory requirements increase quadratically because of the requirement to store sets of pre-calculated inverse-variance matrices. Similarly, for each additional random effect term included in the model, memory requirements increase exponentially. Therefore, we generally limit models to fewer than 10,000 observations and only 1-to-4 random effect terms per trait. There may be opportunities to reduce this memory burden if some of the random effects are low-rank; then these random effects could be updated *on the fly* using efficient routines for low-rank Cholesky updates.

Second, `MegaLMM` is inherently a linear model and cannot effectively model trait relationships that are non-linear. Some non-linear relationships between predictor variables (like genotypes) and traits can be modeled through non-linear kernel matrices, as we demonstrated with the RKHS application to the Bread Wheat data. However, allowing non-linear relationships among traits is currently beyond the capacity of our software and modeling approach. Extending our mixed effect model on the low-dimensional latent factor space to a non-linear modeling structure like a neural network may be an exciting area for future research. Also, some sets of traits may not have low-rank correlation structures that are well-approximated by a factor model. For example, certain auto-regressive dependence structures are low-rank but cannot efficiently be decomposed into a discrete set of factors.

Nevertheless, we believe that in its current form, `MegaLMM` will be useful to a wide range of researchers in quantitative genetics and plant breeding.

### Potential extensions

Beyond the examples we show in this work, the scalability and statistical power of `MegaLMM` can open up new avenues for innovation in genomic prediction applications across the fields of quantitative genetics–both in breeding programs as we have described here and, potentially, in human genetics. Genomic prediction is also used for the calculation of polygenic risk scores for complex human traits and diseases (The International Schizophrenia Consortium 2009). `MegaLMM` may help leverage past case histories, survey responses, molecular tests, and the genetic architecture of other correlated traits to provide a more comprehensive multi-trait polygenic risk score (e.g. Turley *et al.* 2018).

We have focused here on simple scalar phenotypes: the expression of a single gene, the total grain yield, and individual measures of agronomic performance. However, many important traits in plants, animals, and humans cannot easily be reduced to a scalar value. Examples include time-series traits such as growth curves (Campbell *et al.* 2018), metabolic traits such as the relative concentrations of different families of metabolites (Chan *et al.* 2011), and morphological traits such as shape or color (Demmings *et al.* 2019). Each of these traits can be decomposed into vectors of interrelated components, but treating these components as independent prediction targets using existing univariate LMM or low-dimensional MvLMM genomic prediction tools is inefficient because of their statistical dependence. `MegaLMM` can be adapted to make joint predictions on vectors of hundreds or thousands of correlated trait components, which could be fed into high-dimensional selection indices for efficient selection of these important plant characteristics. In human genetics, `MegaLMM` may provide a way to derive multi-ethnic polygenic risk scores (Márquez-Luna *et al.* 2017) by treating outcomes within each ethnic, geographic, or other stratified population group as correlated traits, similar to the analysis of the multi-environment trials above.

`MegaLMM` should be straightforward to extend to more flexible genetic models including the Bayesian Alphabet family of mixture priors on marker effect sizes. These effects can be incorporated into the parameters **B**_2*R*_ and **B**_2*F*_ by adapting the prior structure. This will be further explored in future manuscripts.

Lastly, we have only focused on Gaussian MvLMMs, in which observations are assumed to marginally follow a Gaussian distribution. However, many other types of data require more flexible models. It should be possible to extend `MegaLMM` to the broader family of generalized LMMs. These approaches model the relationships among predictor variables in a latent space, which is then related to the observed data through a link function and an exponential family error distribution. More generally, link-functions could be any non-linear function of multiple parameters such as a polynomial or spline basis, or a mechanistic model. In this case, we would model the correlations among model parameters on this link-scale and then use the link-function to relate the latent scale variables to the observed data. Extending `MegaLMM` to accommodate such generalized LMM structures would require new sampling steps in our MCMC algorithm (see Methods), but we do not see any conceptual challenges with this approach.

## Conclusions

`MegaLMM` is a flexible and powerful framework for the analysis of very high-dimensional datasets in genetics. Multivariate linear mixed models are widely used for analyzing correlated traits, but have been limited to a maximum of a dozen or so traits at a time by the curse of dimensionality. We developed a novel re-parameterization of the MvLMM that allows powerful statistical regularization and efficient computation with thousands of traits. When applied to real plant breeding objectives, `MegaLMM` efficiently leverages information across traits to improve genetic value predictions. Our open-source software package will enable users to apply and extend this method in many directions, opening up new areas of research and development in breeding programs.

## Methods

### Multivariate linear mixed models

Multivariate linear mixed models (MvLMMs) are widely used to model multiple sources of covariance among related observations. Let the *n* × *t* matrix **Y** represent observations on *t* traits for *n* observational units (i.e., individual plants, plots, or replicates).

A general MvLMM takes on the following form

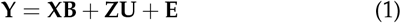

where **X** is a *n* × *b* matrix of “fixed” effect covariates with effect sizes matrix **B**, **U** is an *r* × *t* matrix of random effects for each of the *t* traits, with corresponding random effect design matrix **Z**, and **E** is a *n* × *t* matrix of residuals for each of the *t* traits.

`MegaLMM` uses this formulation to accommodate a large number of designs through different specifications of **X** and **Z**, and different priors on **B**, **U** and **E**. The distinction between “fixed” and “random” effects in Bayesian mixed models is not well-defined because every parameter requires a prior. However, we use the following distinction here: “fixed” effects are covariates assigned flat (i.e., infinite variance) priors or priors with independent variances on each coefficient; “random” effects, in contrast, are grouped in sets that can be thought of as (possibly correlated) samples from a common population distribution. Generally, “fixed” effects are used to model experimental design terms such as blocks, time, sex, etc, genetic principal components, or specific genetic markers; while “random” effects are used to model genetic values, spatial variation, or related effects.

An important feature of `MegaLMM` is that the multiple random effect terms can be included in the model. We specify this as

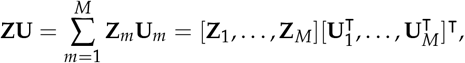

where each **Z**_*m*_ is an *n* × *r_m_* design matrix for a set of related parameters with corresponding coefficient matrix **U**_*m*_. For example, **U**_1_ may model additive genetic values for each individual, while **U**_2_ may model spatial environmental effects for each individual. The distribution of each random effect coefficient matrix is **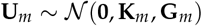**, where 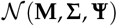 is the matrix normal distribution with mean matrix **M**, among-row covariance **K**_*m*_ and among-column (i.e., among-trait) covariance **G**_*m*_. We assume that both **Z**_*m*_ and **K**_*m*_ are known, while **G**_*m*_ is unknown and must be learned from the data. Note that **K**_*m*_ must be positive semi-definite, while **G**_*m*_ is positive-definite. The covariance among different coefficient matrices is assumed to be zero.

To complete the specification of the MvLMM, we assign the residual matrix the distribution **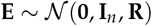** where **I**_*n*_ is the *n* × *n* identity matrix and **R** is an unknown *t* × *t* positive-definite covariance matrix.

### Computational challenges with large multi-trait mixed models

Fitting Eq. (1) is challenging because the columns of **U** and **E** are correlated. This means that data from individual traits (columns of **Y**) cannot be treated independently. Maximum-likelihood approaches for fitting MvLMMs (e.g., `MTG2`) compute the full (or restricted) likelihood of **Y**, which involves calculating the inverse of an *nt* × *nt* matrix many times during model optimization. This is computationally prohibitive when *n* and/or *t* are large (Figure 2A). Gibbs samplers (e.g., `MCMCglmm`) avoid forming and computing the inverse of this extremely large matrix, but still require inverting each of the **G**_*m*_ and **R** matrices repeatedly, which is still prohibitive when *t* is large. Furthermore, the number of parameters in each **G**_*m*_ and **R** grow with the square of *t* and quickly get larger than the total number of observations (*nt*) when *t* is large. This means that **G**_*m*_ and **R** are not identifiable in many datasets and estimates require strong regularization.

### Mixed effect factor model

If both **G**_*m*_ and **R** were diagonal matrices, the *t* traits would be uncorrelated. Fitting Eq. (1) then could be done in parallel across traits, greatly reducing the computational burden. While we cannot directly de-correlate traits, if we can identify the sources of variation that cause trait correlations, the residuals of each trait on these causal factors will be un-correlated. We circumvent this issue by re-parameterizing Eq. (1) as a factor model, where we introduce a set of un-observed (or latent) factors that account for all sources of correlation among the traits. Conditional on the values of these factors, the model reduces to a set of independent linear mixed models. Our re-parameterized multi-trait mixed effect factor model is

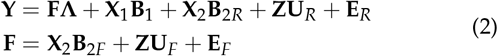

where **F** is an *n* × *K* matrix of latent factors, **Λ** is a *K* × *t* factor loadings matrix, **X**= [**X**_1_, **X**2] is a partition of the *n* × *b* fixed effect covariate matrix between the *b*_1_ covariates with improper priors and the *b*_2_ = *b* − *b*_1_ covariates with proper priors, and **U**_*R*_ and **U**_*F*_ coefficients matrices are specified as:

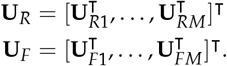

The distributions of the random effects are specified as:

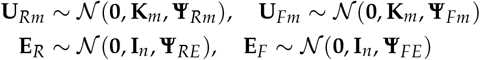

where **Ψ**_*Rm*_, **Ψ**_*Fm*_, **Ψ**_*RE*_, and **Ψ**_*FE*_ are all diagonal matrices. Diagonal elements of **Ψ**_*Fm*_ and **Ψ**_*FE*_ are non-negative, while diagonal elements of **Ψ**_*Rm*_ and **Ψ**_*RE*_ are strictly positive.

Conditional on **F** and **Λ**, the variation in each of the *t* columns of **Y** are uncorrelated and can be fitted to the remaining terms independently. Similarly, the *K* columns of **F** are also uncorrelated and can be modeled independently as well. Therefore, we can fit Eq. (2) without requiring calculating the inverses of any *t × t* matrices, and many calculations can be done in parallel across different CPU cores.

As long as *K* is sufficiently large, Eq. (2) is simply a re-parameterization of Eq. (1). To see how Eq. (2) can represent the terms of Eq. (1), let:

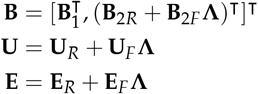

Based on the properties of matrix normal random variables, we can integrate over **U**_*R*_, **U**_*F*_, **E**_*R*_ and **E**_*F*_ to calculate the distributions of each **U**_*m*_ and **E** as:

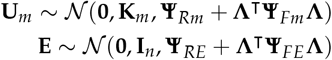

Therefore, each **G**_*m*_ is re-parameterized as **Ψ**_*Rm*_ + **Λ**^⊤^**Ψ**_*Fm*_ **Λ** and **R** is re-parameterized as **Ψ**_*RE*_ + **Λ**^⊤^**Ψ**_*FE*_ **Λ**, such that all off-diagonal elements of each matrix are controlled by **Λ**.

Although these equations appear to imply that our mixed effect factor model constrains **B**, **U** and **E**(and thus each **G**_*m*_ and **R**) to be correlated due to the shared dependence on **Λ**, this is not necessarily the case. When any diagonal element of any **Ψ**_*Fx*_ matrix is set to zero, the corresponding row of **Λ** does not contribute to that term. If at least *t* linearly independent rows of **Λ** contribute to each matrix, then any set of positive-definite matrices can be represented as above. Therefore, we can represent any set of positive-definite matrices **G**_*m*_ and **R** with our model as long as *K >*= *t*(*M* + 1).

Of course, the reason that we parameterize our model in this way is that we do expect some correlation among the genetic and residual covariance matrices. From a statistical perspective, when it is reasonable (given the data) to use the same row of **Λ** for multiple covariance matrices, we can save parameters in the model. From a biological perspective, shared factors provide a biologically realistic explanation for correlations among traits. If we consider the columns of **F** to be *K* traits that simply have not been observed, then it is reasonable to propose that each of these traits is regulated by the same sources of genetic and environmental variation as any of the observed traits.

In Eq. (2), the *K* latent traits (**F**) are the key drivers of all phenotypic co-variation among the *t* observed traits (**Y**). These latent traits may not account for all variation in the observed traits. But, by definition, this residual variation (e.g., measurement errors in each trait) is unique to each trait and uncorrelated with the residual variation in other traits.

#### Prior parameterization

The intuitive structure of the mixed effect factor model (Eq. (2) and Figure 1) makes prior specification and elicitation easier than for Eq. (1) because we do not need to define prior distributions for very large covariance matrices directly. Instead, priors on the random effect variance components and fixed effect regression coefficients are separable and can be described independently, while priors on trait correlations are specified indirectly as priors on the factor loading matrix **Λ**.

In `MegaLMM`, we have chosen functional forms for each prior parameter that balance between interpretability (for accurate prior elicitation), and compatibility with efficient computational approaches. For the variance components, we use the non-parametric discrete prior on variance proportions we previously introduced in `GridLMM` (Runcie and Crawford 2019) that approximates nearly any joint distribution for multiple random effects. For the factor loadings matrix and matrices of regression coefficients, we use a two-dimensional global-local prior based on the horseshoe prior (Carvalho *et al.* 2010), parameterized in terms of the effective number of non-zero coefficients. For the factor loadings matrix specifically, our prior achieves both regularization and interpretability of the factor traits without having to carefully specify *K* itself. Full details of each prior distribution are provided in the Supplemental Methods. Table S1 lists the default hyperparameters for each prior used in the analyses reported here and provided as defaults in the `MegaLMM R` package.

### Computational details and posterior inference

We use a carefully constructed MCMC algorithm to draw samples from the posterior distribution of each model parameter. We gain efficiency in both per-iteration computational time and in effective samples per iteration through careful uses of diagonalization, sparse matrix algebra, parallelization, and integration (or partial collapsing). In particular, our algorithm synthesizes and extends several recent innovations in computational approaches to linear mixed models (Runcie and Mukherjee 2013; Zhou and Stephens 2012; Makalic and Schmidt 2016; Runcie and Crawford 2019). Full details of the computational algorithm are provided in the Supplemental Methods.

### Data Analyses

We demonstrate `MegaLMM` using three example datasets.

#### Scaling performance with gene expression data

To compare the scalability of `MegaLMM` to other multi-trait mixed model programs, we used a large gene expression dataset of 24,175 genes across 728 *Arabidopsis thaliana* accessions. We downloaded the data from NCBI GEO (Barrett *et al.* 2012) (Huang *et al.* GSE80744) and removed genes with average counts *<* 10. We then normalized and variance stabilized the counts using the `varianceStabilizingTransformation` function from `DESeq2` (Love *et al.* 2014). We downloaded a corresponding genomic relationship matrix **K** from the 1001 genomes project (Alonso-Blanco *et al.* 2016) and subsetted to the 665 individuals present in both datasets.

We generated datasets of varying sizes from *t* = 2 to *t* = 24, 175 genes by randomly sampling. We selected one gene as the “focal” trait in each dataset, masked 50% of its values, fit the model in Eq. (1) using four different representative MvLMM programs to the remaining data, and used the results to predict the genetic values of each masked individual for this “focal” gene. Prediction accuracies were estimated as 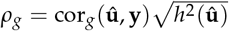, where cor*g* is the estimated genetic correlation evaluated in the testing lines only, and 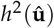 is the heritability of the predictor 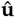 estimated using a univariate LMM (Thompson and Meyer 1986; Lopez-Cruz *et al.* 2020). The simpler Pearson’s correlation estimate of prediction accuracy is not valid in these data because all genes were measured together in the same sample, and therefore some correlation among genes is caused by non-genetic factors (Runcie and Cheng 2019). The four MvLMM prediciton methods were:

1. `MTG2` (Lee and van der Werf 2016): a restricted maximum-likelihood method written in fortran. We pre-calculated the eigenvalue decomposition for **K**, thus this additional time is not included in the results. `MTG2` does not work well with a high percentage of missing data, so genetic value predictions were made with the two-step approach from Runcie and Cheng (2019) which involves estimating model parameters only from the individuals with complete observations, and then incorporating secondary trait values of the new individuals in the second step.
2. `MCMCglmm` (Hadfield 2010): a Bayesian MCMC algorithm largely written in C++. We used “default” priors for **R** and **G** with diagonal means and *ν* = *p*, and ran a single MCMC chain for 7000 iterations, discarding the first 5000 samples as burnin. To speed up calculations (and make the timing results more comparable with the `MegaLMM` algorithm), we rotated the input data by pre-multiplying by the eigenvectors of **K** so that the input relationship matrix was diagonal. Since this matrix rotation is only possible with complete data, we again used the two-step multi-trait prediction approach (Runcie and Cheng 2019).
3. `phenix` (*Dahl et al.* 2016): a variational Bayes algorithm written in `R` that uses a low-rank representation of **G** but a full-rank prior for **R**. We set the maximum number of factors to *p*/4 and used the eigendecomposition of **K** as the input. Again, we excluded this calculation from the time estimates.
4. `MegaLMM`: we ran `MegaLMM` using “default priors” with *K* = min(*n*/4, *p*/2) and collected 6000 MCMC samples, discarding the first 5000 as burnin. We excluded the preparatory calculations, only including the MCMC iterations in the time calculations. For small datasets, these calculations were significant, but were a miniscule part of the analyses of larger datasets.

Each method was run 20 times on different randomly sampled datasets. For the two MCMC methods, we estimated the effective sample size of each element of **U** using the ess_bulk function of the rstan package (Stan Development Team 2019), and used this to estimate the time necessary for the effective sample size to be at least 1000 for 90% of the *u_ij_*. We ran `MTG2` and `MCMCglmm` for datasets up to *t* = 64 because computational times were prohibitively long for larger datasets. We linearly extrapolated the (log) computational times for these methods out to *t* = 512 for comparisons. `phenix` fails when *t* ≥ *n*, so this method is limited to *t <* 665 in this dataset.

To assess the accuracy of each method for estimating genetic and non-genetic covariances, we generated new datasets with 128 genes by calculating empirical correlation matrices for **G** and **R** from two separate samples of 128 genes from the full expression dataset, and then generating genetic and residual values for 128 traits from multivariate normal distributions based on these correlation matrices. For each trait, we converted the correlation matrices into covariance matrices by sampling an independent heritability value for each trait between 0.1 and 0.8. We then estimated the genetic and residual covariance matrices for subsets of these simulated datasets using each of the four above methods. In this example, we found that setting *K* larger (2*p*) gave better results, probably because the **G** and **R** matrices were largely uncorrelated and so independent factors were needed to model the two sets of covariances. Accuracy was reported as the Pearson correlation between the estimated covariance parameters and the true covariance parameters (excluding the variance parameters on the diagonal).

#### Wheat yield prediction using hyperspectral data

We used data from a bread wheat breeding trial to demonstrate how `MegaLMM` can leverage “secondary” traits from high-throughput phenotyping technologies to better predict genetic values of a single target trait. We downloaded grain yield and hyperspectral reflectance data from the bread wheat trials at the Campo Experimental Norman E. Borlaug in Ciudad Obregón, México reported in Krause *et al.* (2019) (Mondal *et al.* 2020). We selected the 2014-2015 Optimal Flat site-year for our main analysis because it had among the greatest number of hyperspectral reflectance data points, and Krause *et al.* (2019) reported relatively low predictive accuracy for grain yield in this site-year. Best linear unbiased estimates (BLUEs) and best linear unbiased predictors (BLUPs) of the line means for grain yield (GY) and 62 hyperspectral bands collected at each of 10 time-points during the growing season, and genotype data from 8519 markers were provided for 1,092 lines in this trial. All other trials were analyzed in the analysis presented in Figure S5.

We compared eight methods for predicting the GY trait based on the genetic marker and hyperspectral data. The first five were “standard” methods using state-of-the-art models for genomic prediction. The final three were new models implemented within the `MegaLMM` framework.

1. `GBLUP`: implemented using the `R` package `rrBLUP` (Endelman 2011), with the genomic relationship matrix **K** calculated with the A.mat function of rrBLUP as in Endelman and Jannink (2012).
2. *Bayesian Lasso* (`BL`): implemented using the `R` package `BGLR` (Perez and de los Campos 2014). We first removed markers with *>* 50% missing data, and imputed the remaining missing genotypes with the population mean allele frequency. We used the default prior parameters for the Bayesian Lasso in BGLR, and collected 9,000 posterior samples with a thinning rate of 5 after a 5,000 iteration burnin.
3. `RKHS`: implemented using `rrBLUP`. We used the same thinned and imputed genotype dataset as for the `BL` method to calculate a genomic distance matrix (**D**). We also used the default `theta.seq` parameter to automatically choose the scale parameter of the Gaussian kernel.
4. `HBLUP`: implemented using the `R` package `lme4qtl`. This replicates the analysis reported by Krause *et al.* (2019), which uses the `GBLUP` method but replaces the genomic relationship matrix described above with **H**, a hyperspectral reflectance relationship matrix calculated as **H**= **SS**^⊤^/620, where **S** is a matrix of centered and standardized BLUEs of hyperspectral bands from each timepoint.
5. `GBLUP+H`: implemented in the `R` package `lme4qtl` (Ziyatdinov *et al.* 2018). This is a two-kernel method, where we use two relationship matrices: **K** and **H**. This method is analogous to the methods proposed by Krause *et al.* (2019) for leveraging the hyperspectral data in prediction; however, those authors only used two-kernel methods for G × E prediction across site-years. Since `lme4qtl` does not predict random effects for un-measured observations, we formed predictions as: 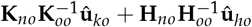 where **K**_*no*_ is the *n_n_* × *n_o_* quadrant of **K** specifying the genomic relationships among the *n_n_* “new” un-observed lines, **K**_*oo*_ is the *n_o_* × *n_o_* quadrant of **K** specifying the genomic relationships among the “old” observed lines, 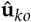 is the vector of BLUPs for the genomic random effect, and **H**_*no*_, **H**_*oo*_ and 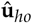 are similar quantities for the hyperspectral random effect.
6. `MegaLMM-GBLUP`: we modeled the combined trait data **Y**= [**y**, **S**] with the model specified in Eq. (2) using a single random effect with relationship matrix **K** as above, no fixed effects besides an intercept (**X** was a column of ones and **X**2 had zero columns). We ran `MegaLMM` with *K* = 100 factors, “default” priors (see Table S1), and two partitions of the trait data (the first containing grain yield with the masked training set as described below, and the second containing all 620 hyperspectral bands with complete data). We collected 500 posterior samples of the quantity: **u**_1_ = **u**_*R*1_ + (**U***F **λ***_1_) at a thinning rate of 2, discarding the first 1,000 iterations as burn-in.
7. `MegaLMM-RKHS`: we implemented multi-trait RKHS regression model using the “kernel-averaging” method proposed by de Los Campos *et al.* (2010). We standardized **D** based on its mean (squared) value, and placed a uniform prior on the set of scaling factors *h* = {1/5, 1, 5}, which we implemented by calculating three corresponding relationship matrices **K**_1_,…, **K**_3_ and by specifying three random effects in Eq. (2). We again used “default” priors, *K* = 100 factors, and treated only the global intercept per-trait as fixed effects. We collected 500 posterior samples of the quantity: **Zu**_1_ = **Zu**_*R*1_ + **Z**(**U***_F_ **λ***_1_) at a thinning rate of 2, discarding the first 1000 iterations as burn-in.
8. `MegaLMM-GBLUP-CV1`: we repeated the `MegaLMM-GBLUP` method above, but this time without partitioning the trait data. Instead, we masked both the grain yield and the 620 hyperspectral band data from the testing set so all lines in the training data had complete data. Predictions of the genetic values were calculated identically to above.

We used cross-validation to evaluate the prediction accuracy of each method. We randomly selected 50% of the lines for model training, 50% for testing, and masked the GY observations for the testing lines. We fit each model to the partially-masked dataset and collected the predictions of GY for the testing lines. We estimated prediction accuracy as 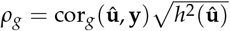 because the hyperspectral reflectance data were collected on the same plots as the GY data and therefore non-genetic (i.e., microenvironmental) factors that affect both reflectance and yield may induce non-genetic correlations among traits (Runcie and Cheng 2019). BLUPs were used as the predictand except in the 2016-17 year when the BLUPs were poorly corelated with the BLUEs suggesing data quality issues. We used a 50-50 training-testing split of the data to ensure that cor*_g_* could be estimated accurately in the testing partition. This cross-validation algorithm was repeated 20 times with different random partitions.

#### Maize trait imputation in multi-environment trails

We used data on maize hybrids from the Genomes-To-Fields Initiative experiments to demonstrate how `MegaLMM` can leverage genetic correlations across locations in multi-environment trials. We downloaded the agronomic data from the 2014-2017 field seasons from the CyVerse data repository (McFarland *et al.* 2020) and corresponding genomic data. We used TASSEL5 (Bradbury *et al.* 2007) to build a kinship matrix for each hybrid genotype using the CenteredIBS routine.

A total of 2012 non-check hybrids with phenotype and genotype data from 108 trials (i.e., site-years) were available. We selected four representative agronomic traits: plant height (cm), grain yield (bushels/acre), days-to-silking (days), and the anthesis-silking interval (ASI, days). For each trait in each site-year, we calculated BLUPs for all observed genotypes using the `R` package `lme4` (Bates *et al.* 2015) with Rep and Block:Rep as fixed effects to account for the experimental design in each field, and formed them into 2012 × 108 BLUP matrices for each trait. We then dropped site-years where the BLUP variance was zero, or which had fewer than 50 tested lines. On average ≈ 12% of hybrid-site-year combinations were observed across each of the four BLUP matrices. We then used four methods to predict the BLUPs of hybrids that were not grown in each trial:

1. `GBLUP (univariate)`: missing values were imputed separately for each site:year using the `mixed.solve` function of the `rrBLUP` package.
2. `GBLUP (env BLUPs)`: genetic values for each hybrid were assumed to be constant across all site-years. We estimated these in two steps. In the first step, we estimated hybrid main effects treating lines as independent random effects using `lme4`, with `site:year` included as a fixed effect. In the second step, we estimated genetic values using the `mixed.solve` function of the `rrBLUP` package.
3. phenix: we used phenix to impute missing observations in **Y** using **K** as a relationship matrix.
4. `MegaLMM`: we fit the model specified in Eq. (2) to the full matrix **Y**, with *K* = 50 factors and “default”. Here, we partitioned **Y** into 4 sets based on year to minimize the number of missing observations to condition on during the MCMC. We collected 1000 posterior samples of imputed values 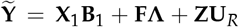 with a thinning rate of 2, after discarding the first 5000 iterations as burnin.

We estimated prediction accuracy of each method using cross-validation. For each of 20 replicate cross-validation runs per model, we randomly masked 20% of the non-missing BLUPs, and then calculated the Pearson’s correlation between these “observed” values and the imputed values of each method. Pearson’s correlation is appropriate as an estimate of genomic prediction accuracy in this case because different plants were used in each trial, so there is no non-genetic source of correlation among site-years that may bias accuracy estimates.

## Declarations

### Ethics approval and consent to participate

Not applicable

### Consent for publication

Not applicable

### Availability of data and materials

All data used in these analyses were downloaded from the publicly accessible repositories described above. Arabidopsis gene expression data was downloaded from the NCBI GEO accession GSE80744 available at https://www.ncbi.nlm.nih.gov/geo/query/acc.cgi?acc=GSE80744.

The Arabidopsis kinship matrix was downloaded from https://1001genomes.org/data/GMI-MPI/releases/v3.1/SNP_matrix_imputed_hdf5/1001_SNP_MATRIX.tar.gz. The wheat dataset was downloaded from the CIMMYT Research Data & Software Repository Network available at http://hdl.handle.net/11529/10548109. The maize phenotype data were downloaded from the CyVerse data repository based on the links described in (*McFarland et al.* 2020). Genomic data were downloaded from (http://datacommons.cyverse.org/browse/iplant/home/shared/commons_repo/curated/Carolyn_Lawrence_Dill_G2F_Nov_2016_V.3/b._2014_gbs_data). Scripts for running analyses are available in the GitHub repository: https://github.com/deruncie/MegaLMM_analyses. The `R` package for `MegaLMM` is available here: https://github.com/deruncie/MegaLMM/tree/v0.9.1 and is licensed with the Polyform Noncommercial 1.0 license. The specific versions of the scripts and package codes are archived at zenodo with DOIs 10.5281/zenodo.4735048 and 10.5281/zenodo.4740662.

### Competing interests

The authors declare that they have no competing interests

### Funding

This work is supported by Agriculture and Food Research Initiative grants no. 2020-67013-30904 and 2018-67015-27957 from the USDA National Institute of Food and Agriculture to DER and HC. DER is also supported by United States Department of Agriculture (USDA) National Institute of Food and Agriculture (NIFA), Hatch project 1010469. LC is supported by grants P20GM109035 (COBRE Center for Computational Biology of Human Disease; PI Rand) and P20GM103645 (COBRE Center for Central Nervous; PI Sanes) from the NIH NIGMS, 2U10CA180794-06 from the NIH NCI and the Dana Farber Cancer Institute (PIs Gray and Gatsonis), as well as by an Alfred P. Sloan Research Fellowship and a David & Lucile Packard Fellowship for Science and Engineering.

### Authors’ contributions

DER developed the method, wrote the `R` package, developed and ran the analyses, and wrote the paper. JQ edited the manuscript HC helped develop the method, design the analysis, and edited the paper LC helped develop the method, design the analysis, and wrote the paper

## Acknowledgements

Not applicable

## Notes

https://github.com/deruncie/MegaLMM

https://github.com/deruncie/MegaLMM_analyses

### Competing Interest Statement

The authors have declared no competing interest.

### Summary of Updates

updated text

