## Supplemental Methods and Figures for "MegaLMM: Mega-scale linear mixed models for genomic predictions with thousands of traits"

### 1 Supplementary Methods

#### 1.1 Prior Parameterization

In the `MegaLMM` software, we have chosen functional forms for each prior parameter that balance between interpretability (for accurate prior elicitation), and compatibility with efficient computational algorithms. We detail these choices below.

**Variance Components.** The MegaLMM model (see Eq. (2) in main text) has $(t+K)(M+1)$ variance component parameters that need to be estimated (i.e., diagonal elements of the $\mathbf{\Psi}$ matrices). Most Bayesian linear mixed models (LMMs) place independent inverse-Gamma priors on each variance component. This is a convenient choice due to conjugacy with a Gaussian likelihood. However, inverse-Gamma priors can cause problems with mixing in Gibbs samplers (particularly when the variance component is close to zero) (Gelman 2006). Default hyperparameters for inverse-Gaussian distributions also lead to non-intuitive surfaces for the joint distribution of two or more variance components, which favor models where one random effect dominates over the others (Runcie and Crawford 2019). In our experience, eliciting priors for the *proportion of variance* attributable to each random effect is more intuitive than eliciting priors for the absolute value of each variance. Below, we use the symbol $h_m^2$ to represent the proportion of total variance attributable to random effect $m$ because of the parallel between this term and the concept of heritability.

For each of the $t$ observed traits, let the variance parameters be denoted by $\boldsymbol{\psi}_{Rj} = (\psi_{R1j}, \ldots, \psi_{RMj}, \psi_{REj})$. We specify priors on these parameters indirectly by re-parameterizing them as

$$\boldsymbol{\psi}_{Rj} = \sigma_{Rj}^2 \mathbf{h}_{Rj}^2, \tag{S1}$$

where $\sigma_{Rj}^2 = \sum_m \boldsymbol{\psi}_{Rmj} + \psi_{REj}$ is used to denote the total variance and $\mathbf{h}_{Rj}^2 = (h_{R1j}^2, \ldots, h_{RMj}^2, h_{REj}^2)$ with each individual proportion being equal to $h_{Rmj}^2 = \psi_{Rmj}/\sigma_{Rj}^2$. We do end up using an inverse-gamma for the prior distribution on the total variance term $\sigma_{Rj}^2$ because estimates are generally not near zero (unless all variation in a trait is explained by $\mathbf{X}$ or $\mathbf{F}\boldsymbol{\lambda}_j$) and it is the only variance parameter for each trait in the re-parameterized model.

We allow virtually any prior specification for $\mathbf{h}_{Rj}^2$ (including non-parametric distributions) on the $M$-dimensional simplex (such that the vector sums to one). We do this by approximating the prior surface over a pre-defined grid of points. This follows our earlier work with single-trait linear mixed models (Runcie and Crawford 2019) where we showed that such grid-interpolations can be leveraged for massive computational gains without appreciable loss of accuracy in estimating moments of posterior distributions. In the proceeding sections, we describe how the Gibbs sampler implemented in `MegaLMM` takes advantage of this discretized prior surface for improved MCMC sampling and faster computation.

For the $K$ latent factor traits, the variance parameters are denoted by $\boldsymbol{\psi}_{Fk} = (\psi_{F1k}, \ldots, \psi_{FMk}, \psi_{FEk})$, which we again re-parameterize as $\boldsymbol{\psi}_{Fk} = \sigma_{Fk}^2 \mathbf{h}_{Fk}^2$. However, to ensure identifiability, we set $\sigma_{Fk}^2 = 1$. We then allow the same discretized prior to be used for $\mathbf{h}_{Fk}^2$ as we just described.

**Factor Loadings Matrix.** Rows of $\mathbf{\Lambda}$ hold the regression coefficients that describe the relationship between the observed and latent factor traits. With $K$ factors and $t$ traits, there are $Kt$ regression coefficients, so strong regularization is required when $t$ and/or $K$ is large. We use a two-dimensional global-local shrinkage approach based on the horseshoe prior to achieve both regularization and interpretability on the factor traits without having to carefully specify $k$ itself. Within each row of $\mathbf{\Lambda}$, the local shrinkage part of the prior pushes the unimportant coefficients strongly towards zero. This step is exchangeable across traits, reflecting a lack of information on which traits may be correlated. Global shrinkage is induced across rows, where regularization on all coefficients for the $(k+1)$-th latent trait is done relative to the $k$-th. Here, we draw on the "sparse infinite factor" prior from Bhattacharya and

Dunson (2011) which induces ordering of the latent traits from the most-to-least important by requiring that the magnitude of the global shrinkage parameter stochastically increases from one latent trait to the next. This enforces that the expected number of nonzero elements in row $k+1$ is smaller than the number of nonzero elements in row $k$. This means that high-order factors are less important, and we can choose a threshold beyond which we can safely ignore any higher-order factors.

We parameterize this two-dimensional prior in terms of the expected proportion of approximately zero regression coefficients in the first (i.e., most important) factor, and the expected change in this proportion as we move from from factor $k$ to factor $k+1$. Our prior for $\mathbf{\Lambda}$ has the following form:

$$\begin{aligned} \lambda_{kj} \mid \sigma^2_{Rj} &\sim \mathcal{N}(0, \phi^2_{kj}\tau^2_k\sigma^2_{Rj}), \\ \tau^2_k &= \tau^2 \prod_{h=1}^{k} \delta_h, \\ \phi_{kj} &\sim \mathcal{C}^+(0,1) \\ \tau &\sim \mathcal{C}^+(0,\tau_0) \\ \delta_h &\sim iG(a_\delta, b_\delta), \end{aligned} \tag{S2}$$

where we fix $\delta_1 = 1$, $\mathcal{C}^+(0,s)$ is the half-Cauchy distribution with scale $s$, $\sim iG(a,b)$ is the inverse Gamma distribution with shape $a$ and scale $b$, and $\phi_{kj}$ provides the local-shrinkage on each $\lambda_{kj}$, while $\tau_k$ provides global-shrinkage for the $k$-th row of $\mathbf{\Lambda}$ (which we denote as $(\mathbf{\Lambda}^\intercal)_k$).

To choose hyperparameters $\tau_0$, $a_\delta$ and $b_\delta$, we consider how each controls the expectation of the number of effectively nonzero coefficients in each row of $\mathbf{\Lambda}$ (no parameter will be exactly zero but the horseshoe prior shrinks non-informative priors close to zero). Using the approximate shrinkage-factor calculation from Piironen and Vehtari (2017) and assuming that columns of $\mathbf{F}$ have unit variance, we can estimate the effective number of nonzero coefficients in row $k$ given the global shrinkage factor $\tau_k$ as

$$m_k \mid \tau_k = \sum_{j=1}^{t} (1-\kappa_{kj}) \tag{S3}$$

where $\kappa_{kj} = (1+n\phi^2_{kj}\tau^2_k)^{-1}$ is the shrinkage factor for each coefficient. Note: this equation differs slightly from equation 2.4 in (Piironen and Vehtari 2017) because in their formulation the prior variance of the regression coefficients is not proportional to $\sigma^2$ (equation 2.2). Therefore, the $\sigma^{-2}$ term cancels in our calculation of $\kappa_{kj}$. Piironen and Vehtari (2017) showed that the expectation of $m_k \mid \tau_k$ simplifies to

$$\mathrm{E}[m_k \mid \tau_k] = \frac{\tau_k\sqrt{n}}{1+\tau_k\sqrt{n}} t. \tag{S4}$$

Thus, the quantity $\tau_k\sqrt{n}$ can be interpreted as the odds of each coefficient in $(\mathbf{\Lambda}^\intercal)_k$ being nonzero. If we knew that there were exactly $t^*_k$ non-zero coefficients of $(\mathbf{\Lambda}^\intercal)_k$, we could solve for $\tau_k = t^*_k/(\sqrt{n}(t-t^*_k))$.

We start with a prior guess of $t_0$ non-zero regression coefficients for the first and most-important factor $(\mathbf{\Lambda}^\intercal)_1$. This means that $\tau_1 = \tau$ since $\delta_1 = 1$, and so we set the hyperparameter $\tau_0$ to the value $t_0/(\sqrt{n}(t-t_0))$. Since $\tau \sim \mathcal{C}^+(0,\tau_0)$, the median of the prior distribution for the quantity $\tau_1\sqrt{n}$ will be $t_0/(t-t_0)$. Next, for $(\mathbf{\Lambda}^\intercal)_2$, $\tau_2 = \tau_1\sqrt{\delta_2}$, so the median of $\tau_2\sqrt{n}$ (i.e., the prior guess for $t_2/(t-t_2)$) will be $\sqrt{\delta_2}t_0/(t-t_0)$–which is $\sqrt{\delta_2}$ times the odds for each coefficient of $\boldsymbol{\lambda}_1$. The same pattern will repeat for each succeeding factor, with its odds for each coefficient being distinguishable from zero changing by a factor of $\sqrt{\delta_k}$ relative to the previous factor. As long as $\mathrm{E}[\delta_k] < 1$, these odds will decrease for higher-order factors, eventually reaching close to zero. We use this to calibrate the hyperparameters $a_\delta$ and $b_\delta$, either by simulating from the prior or by using the approximation that $\mathrm{E}[\sqrt{1/\delta_k}] \approx \sqrt{a_\delta/b_\delta}$. For example, setting $a_\delta = 3$ and $b_\delta = 1$ gives a mean decrease in odds of 1.7× and a 91% probability of the change in odds being between 1× and 3×.

It is important to note that we do not try to learn the number of factors $K$. The advantage of the "infinite factor model" prior is that higher-rank factors are shrunk strongly towards zero in a data-dependent way. Therefore, the actual value of $K$ tends to be unimportant as long as it is large enough such that all important factors can be included. Making $K$ larger than that value has little impact on the posterior predictions or inferences of other parameters. We can thus save computation by pruning factors with all small coefficients.

Note that our prior here differs from the priors proposed by Bhattacharya and Dunson (2011) and Runcie and Mukherjee (2013) because of our decision to use a half-Cauchy distribution for the local-shrinkage parameters instead of the inverse-gamma distribution. The inverse-gamma distribution induces a t-distribution on the regression coefficients. While this prior generally performs well, and we include it as an option in our *R* package, we find that eliciting priors for the local and global shrinkage with this distribution is not intuitive. The horseshoe prior is generally better at separating signal from noise, shrinking only unimportant coefficients closer to zero—and thus leads to more interpretable factors.

**Fixed Effects.** As noted above, we partition the covariates in $\mathbf{X}$ into the $b_1$ covariates with improper priors on their coefficients, and the remaining $b_2 = b - b_1$ covariates with proper priors on their coefficients. $\mathbf{B}_1$ is the matrix of regression coefficients for the covariates with improper priors, where we assume each element $b_{bj} \sim \mathcal{N}(0, \infty)$.

When included, covariates in $\mathbf{X}_2$ are generally high-dimensional (e.g., genetic markers), often with $b_2 > n$. Therefore, we use priors that regularize estimates of the coefficients $\mathbf{B}_{2R}$ and $\mathbf{B}_{2F}$ and favor sparse, interpretable solutions. $\mathbf{B}_{2R}$ plays a very similar role to the factor loadings matrix $\mathbf{\Lambda}$ in `MegaLMM` (see Eq. (2) in the main text), as the model involves the combined effect: $\mathbf{X}_2\mathbf{B}_{2R} + \mathbf{F\Lambda} = [\mathbf{X}_2, \mathbf{F}][\mathbf{B}_{2R}^\intercal, \mathbf{\Lambda}^\intercal]^\intercal$ where $\mathbf{X}_2$ and $\mathbf{F}$ are both covariate matrices (with $\mathbf{X}_2$ known *a priori* and $\mathbf{F}$ unknown), and $\mathbf{B}_{2R}$ and $\mathbf{\Lambda}$ are the corresponding coefficient matrices. Due to this connection, our default prior for $\mathbf{B}_{2R}$ is also the horseshoe prior. However in this case we apply independent global shrinkage parameters for each trait $j$ (i.e., columns of $\mathbf{B}_{2R}$).

We also use a horseshoe prior for each column of $\mathbf{B}_{2F}$. A key feature of `MegaLMM` is that we split the total effect of covariate $\mathbf{x}_{2l}$ into two components: $\mathbf{x}_{2l}(\mathbf{B}_{2R}^\intercal)_l + \mathbf{x}_{2l}(\mathbf{B}_{2F}^\intercal)_l\mathbf{\Lambda}$, such that the $K$-dimensional row-vector $(\mathbf{B}_{2F}^\intercal)_l$ partially accounts for the effects of $\mathbf{x}_{2l}$ through the common factors, and the $t$-dimensional vector $(\mathbf{B}_{2R}^\intercal)_l$ accounts for the remain effects. Without regularization on $\mathbf{B}_{2F}$ and $\mathbf{B}_{2R}$, either would provide equivalent explanations for the data and the two would not be simultaneously identifiable. However, when we can explain the effects of $\mathbf{x}_{2l}$ on $\mathbf{Y}$ effectively with the $K$ values in $(\mathbf{B}_{2F}^\intercal)_l$ (conditional on $\mathbf{\Lambda}$) as we can do with the $t$ values of $(\mathbf{B}_{2R}^\intercal)_l$, our prior favors the former solution as it is sparser and we end up with a more parsimonious model.

#### 1.2 MCMC Algorithm

Our hybrid Gibbs and Metropolis-Hastings sampler uses the following steps:

1. *Sample* $[\mathbf{B}_1, \mathbf{B}_{2R}, \mathbf{\Lambda}, \mathbf{U}_{Rm}, \mathbf{\Phi}_{\mathbf{b}_{2R}}, \mathbf{\Phi}_{\Lambda}, \boldsymbol{\tau}, \boldsymbol{\tau}_{\mathbf{b}_{2R}}, \mathbf{h}^2_{Rj}, \sigma^2_{Rj}]$ *given* $[\mathbf{Y}, \mathbf{F}]$. The priors for $\mathbf{B}_1$, $\mathbf{B}_{2R}$, $\mathbf{\Lambda}$ and $\mathbf{U}_{Rm}$ factorize into independent distributions for each of their $t$ columns (conditional on $\boldsymbol{\delta}$). The priors for $\mathbf{h}^2_{Rj}$ and $\sigma^2_{Rj}$ are also independent for $j = 1 \dots t$. Therefore, conditional on $\mathbf{F}$ (and $\boldsymbol{\delta}$), we can simplify Eq. (2) in the main text into $t$ independent univariate linear mixed models of the form:

$$
\begin{aligned}
\widetilde{\mathbf{y}}_j &= \widetilde{\mathbf{X}}_1\mathbf{b}_{1j} + \widetilde{\mathbf{F}}\boldsymbol{\lambda}_j + \widetilde{\mathbf{X}}_2\mathbf{b}_{2Rj} + \widetilde{\mathbf{Z}}\mathbf{u}_{Rj} + \widetilde{\mathbf{e}}_{Rj} \\
\mathbf{b}_{1j} &\sim \mathcal{N}(\mathbf{0}, \infty\mathbf{I}_{b_1}) \\
\mathbf{b}_{2Rj} &\sim \mathcal{N}(\mathbf{0}, \sigma^2_{Rj}\mathbf{D}_{\mathbf{b}_{2Rj}}) \\
\boldsymbol{\lambda}_j &\sim \mathcal{N}(\mathbf{0}, \sigma^2_{Rj}\mathbf{D}_{\lambda_j})
\end{aligned}
$$

$$\mathbf{u}_{Rj} \sim \mathcal{N}(\mathbf{0}, \sigma^2_{Rj}\mathbf{K}(\mathbf{h}^2_{Rj}))$$
$$\widetilde{\mathbf{e}}_{Rj} \sim \mathcal{N}(\mathbf{0}, \sigma^2_{Rj} h^2_{REj}\mathbf{I}_{\widetilde{n}_j})$$
$$\sigma^2_{Rj} \sim \mathrm{iG}(a_R, b_R),$$

where $\mathbf{y}_j$ is the $j$th column of $\mathbf{Y}$, $\boldsymbol{\lambda}_j$ is the $j$th column of $\boldsymbol{\Lambda}$, $\widetilde{\mathbf{y}}_j$ denotes the elements of $\mathbf{y}_j$ that are non-missing (similar definitions are used for the tildes over other matrix and vector terms), $\mathbf{D}_{\mathbf{b}_{2Rj}} = \mathrm{diag}(\tau^2_{\mathbf{b}_{2R}j}\boldsymbol{\phi}^2_{\mathbf{b}_{2R}\cdot j})$ and $\mathbf{D}_{\lambda_j} = \mathrm{diag}(\boldsymbol{\phi}^2_{\cdot j} \odot \boldsymbol{\tau}^2)$ are diagonal matrices composed of the prior variances for each element of $\mathbf{b}_{2Rj}$ or $\boldsymbol{\lambda}_j$ (excluding the term $\sigma^2_{Rj}$), and $\mathbf{K}(\mathbf{h}^2)$ is a matrix-valued function returning an $r \times r$ matrix as a function of $\mathbf{h}^2$, generally a block-diagonal matrix with each block the product of an element of $\mathbf{h}^2$ and a pre-defined covariance matrix $K_m$. Only non-missing elements $\widetilde{\mathbf{y}}_j$ contribute to the likelihood, so the remainder of $\mathbf{y}_j$ can be ignored.

We collect samples of the coefficients $\{\mathbf{b}_{1j}, \mathbf{b}_{2Rj}, \boldsymbol{\lambda}_j, \mathbf{u}_{Rj}\}$ and variance parameters $\{\mathbf{D}_{\mathbf{b}_{2Rj}}, \mathbf{D}_{\lambda_j}, \mathbf{h}^2_{Rj}, \sigma^2_{Rj}\}$ using a set of highly optimized Gibbs-type updates based on the fact that many of the same intermediate calculations can be re-used among different columns of $\mathbf{Y}$. Full derivations and sampling distributions of each step are described below in Supplementary Section 1.3.

2. *Sample* $[\mathbf{B}_{2F}, \mathbf{U}_{Fm}, \boldsymbol{\Phi}_{\mathbf{b}_{2F}}, \boldsymbol{\tau}_{\mathbf{b}_{2F}}, \mathbf{h}^2_F]$ *given* $\mathbf{F}$. Given the parallelism between the models for $\mathbf{F}$ and $\mathbf{Y}$, the sampling steps for the parameters of the $\mathbf{F}$ model are analogous to those described above. Again, the models for the $k$ columns of $\mathbf{F}$ are independent and each has the form:

$$\mathbf{f}_k = \mathbf{X}_2\mathbf{b}_{2Fk} + \mathbf{Z}\mathbf{u}_{Fk} + \mathbf{e}_{Fk}$$
$$\mathbf{b}_{2Fk} \sim \mathcal{N}(\mathbf{0}, \mathbf{D}_{\mathbf{b}_{2Fk}})$$
$$\mathbf{u}_{Fk} \sim \mathcal{N}(\mathbf{0}, \mathbf{K}(\mathbf{h}^2_{Fk}))$$
$$\mathbf{e}_{Fk} \sim \mathcal{N}(\mathbf{0}, h^2_{FEk}\mathbf{I}_n),$$

where $\mathbf{D}_{\mathbf{b}_{2Fk}} = \mathrm{diag}(\tau^2_{\mathbf{b}_{2F}k}\boldsymbol{\phi}^2_{\mathbf{b}_{2F}\cdot k})$. The main difference from above is the lack of the term $\sigma^2_{Fk}$ in each prior distribution. This is missing because $\sigma^2_{Fk} = 1$ for identifiability.

3. *Sample* $\mathbf{F}$ *given all other parameters.* To sample $\mathbf{F}$, we transpose Eq. (2) in the main text to:

$$\mathbf{Y}^\intercal = \boldsymbol{\Lambda}^\intercal\mathbf{F}^\intercal + \mathbf{M}^\intercal_R + \mathbf{E}^\intercal_R$$

where $\mathbf{M}_R = \mathbf{X}_1\mathbf{B}_1 + \mathbf{X}_2\mathbf{B}_{2R} + \mathbf{Z}\mathbf{U}_R$ and $\mathbf{F}$ is simply a set of linear regression coefficients. Furthermore, by conditioning on $\mathbf{B}_{2R}$, $\mathbf{U}_R$, $\mathbf{B}_{2F}$ and $\mathbf{U}_F$, columns of $\mathbf{F}^\intercal$ and $\mathbf{M}^\intercal_R$ are uncorrelated and we can represent this as a set of $n$ simple linear regressions:

$$(\widetilde{\mathbf{Y}}^\intercal)_i = \widetilde{\boldsymbol{\Lambda}}^\intercal(\mathbf{F}^\intercal)_i + (\widetilde{\mathbf{M}}^T_R)_i + (\widetilde{\mathbf{E}}^\intercal_R)_i$$
$$(\mathbf{F}^\intercal)_i \sim \mathcal{N}(\boldsymbol{\mu}_{(\mathbf{F}^\intercal)_i}, \mathbf{D}_\mathbf{f})$$
$$(\widetilde{\mathbf{E}}^\intercal_R)_i \sim \mathcal{N}\Big(\mathbf{0}, \mathbf{D}_{(\widetilde{\mathbf{Y}}^\intercal)_i}\Big)$$

where $(\widetilde{\mathbf{Y}}^\intercal)_i$ is the sub-vector composed of the non-missing traits in the $i$th row of $\mathbf{Y}$, $\boldsymbol{\mu}_{(\mathbf{F}^\intercal)_i} = \mathbf{B}^\intercal_{2F}(\mathbf{X}^\intercal_2)_i + \mathbf{U}^\intercal_F(\mathbf{Z}^\intercal)_i$, $\mathbf{D}_\mathbf{f} = \mathrm{diag}(h^2_{FE_k})$ is a diagonal matrix holding the residual variances of each of the $K$ columns $\mathbf{f}_k$, and $\mathbf{D}_{(\widetilde{\mathbf{Y}}^\intercal)_i} = \mathrm{diag}(h^2_{RE_j}\sigma^2_{Rj})$ is the similar diagonal matrix with the residual variances of $(\widetilde{\mathbf{Y}}^\intercal)_i$. Therefore, the conditional posterior is $(\mathbf{F}^\intercal)_i \,|\, \cdot \sim \mathcal{N}(\boldsymbol{\mu}, \boldsymbol{\Sigma})$ where

$$\boldsymbol{\Sigma} = \left[\boldsymbol{\Lambda}\mathbf{D}^{-1}_{(\widetilde{\mathbf{Y}}^\intercal)_i}\boldsymbol{\Lambda}^\intercal + \mathbf{D}^{-1}_\mathbf{f}\right]^{-1}$$
$$\boldsymbol{\mu} = \boldsymbol{\Sigma}\left[\boldsymbol{\Lambda}\mathbf{D}^{-1}_{(\widetilde{\mathbf{Y}}^\intercal)_i}\Big((\widetilde{\mathbf{Y}}^\intercal)_i - (\widetilde{\mathbf{M}}^T_R)_i\Big) + \mathbf{D}^{-1}_\mathbf{f}\boldsymbol{\mu}_{(\mathbf{F}^\intercal)_i}\right].$$

In the case where we wish to impute $(\mathbf{F}^\intercal)_i^*$ for an individual with no observations (i.e., when $(\widetilde{\mathbf{Y}}^\intercal)_i^*$ is empty), we simply draw $(\mathbf{F}^\intercal)_i^* \sim \mathcal{N}(\boldsymbol{\mu}_{(\mathbf{F}^\intercal)_i}, \mathbf{D_f})$.

4. *Sample missing elements of* $\mathbf{Y}$ *given all other parameters.* Although missing observations are generally not needed for sampling the model parameters (except in cases described below), imputing them is useful for predicting unmeasured phenotypes. The conditional posterior of each element follows a univariate Gaussian distribution in which

$$y_{ij} \sim \mathcal{N}(\mu_{ij}, h^2_{REj}\sigma^2_{Rj}),$$

where $\mathbf{M} = \{\mu_{ij}\} = \mathbf{F\Lambda} + \mathbf{X}_1\mathbf{B}_1 + \mathbf{X}_2\mathbf{B}_{2R} + \mathbf{ZU}_R$.

#### 1.3 Gibbs Sampler Updates

We now detail specific Gibbs sampler updates used in `MegaLMM`. Steps 1 and 2 above both involve sets of parallel linear regression models. Although the design matrices may differ for columns of $\mathbf{Y}$ and $\mathbf{F}$, the form of both sets of conditional models (replacing specific variable names and dropping subscripts) is:

$$\begin{aligned}
\mathbf{y} &= \mathbf{X}_1\boldsymbol{\alpha} + \mathbf{X}_2\boldsymbol{\beta} + \mathbf{Zu} + \mathbf{e} \\
\boldsymbol{\alpha} &\sim \mathrm{N}(\mathbf{0}, \infty) \\
\boldsymbol{\beta} &\sim \mathrm{N}(\mathbf{0}, \sigma^2\mathbf{D}_\beta) \\
\mathbf{u} &\sim \mathrm{N}(\mathbf{0}, \sigma^2\mathbf{K}(\mathbf{h}^2)) \\
\mathbf{e} &\sim \mathrm{N}(\mathbf{0}, \sigma^2 h_e^2\mathbf{I}) \\
\sigma^2 &\sim \mathrm{iG}(a_0, b_0)
\end{aligned} \tag{S5}$$

where $\mathbf{y}$ is an $n$-dimensional vector of observations; $\{\boldsymbol{\alpha}, \boldsymbol{\beta}, \mathbf{u}\}$ are vectors of unknown coefficients of length $a, b$ and $r$, respectively, with known design matrices of appropriate size; and $\mathbf{e}$ is an $n$-dimensional vector of independent and identically distributed residuals. We require $\mathbf{D}_\beta$ to be diagonal, while $\mathbf{K}(\mathbf{h}^2)$ is a matrix-valued function returning an $r \times r$ matrix as a function of $\mathbf{h}^2$. In Step 1: $\mathbf{y} = \widetilde{\mathbf{y}}_j$, $\boldsymbol{\alpha} = \mathbf{b}_{1j}$, $\boldsymbol{\beta} = [\boldsymbol{\lambda}_j^\intercal, \mathbf{b}_{2Rj}^\intercal]^\intercal$, $\mathbf{e} = \widetilde{\mathbf{e}}_{Rj}$, and $\sigma^2 = \sigma^2_{Rj}$. In Step 2, $\mathbf{y} = \mathbf{f}_k$, $\boldsymbol{\alpha}$ is empty, $\boldsymbol{\beta} = \mathbf{b}_{2Fk}$, $\mathbf{e} = \mathbf{e}_{Fk}$, and $\boldsymbol{\sigma}^2 = 1$. Lastly, we use $\mathrm{iG}(\alpha, \beta)$ to denote the inverse-Gamma distribution with probability density function

$$p(z|\alpha, \beta) = \frac{\beta^\alpha}{\Gamma(\alpha)} z^{-\alpha-1} \exp\left\{-\frac{\beta}{z}\right\}.$$

Our goal is to draw samples for all unknown parameters as efficiently as possible while minimizing autocorrelation in the MCMC chain. We accomplish this by blocking (or collapsing) many sampling steps where we first integrate out certain regression coefficients and then draw samples for these in subsequent steps. Note that, in a single iteration, we draw samples for all parameters for many (say $p$) $\mathbf{y}$ vectors. Since the parameters $\{\mathbf{X}_1, \mathbf{X}_2, \mathbf{Z}, \mathbf{K}(\cdot)\}$ are constant for each column of $\mathbf{Y}$ or $\mathbf{F}$, we cache as many of the intermediate calculations as possible to reduce unnecessary operations. Our sampling strategy has the following steps:

1. Sample $\boldsymbol{\alpha} \,|\, \mathbf{D}_\beta, \mathbf{h}^2, h_e^2, \sigma^2$, integrating out $\boldsymbol{\beta}, \mathbf{u}$.
2. Sample $\sigma^2 \,|\, \boldsymbol{\alpha}, \mathbf{D}_\beta, \mathbf{h}^2, h_e^2$, integrating out $\boldsymbol{\beta}, \mathbf{u}$.
3. Sample $\boldsymbol{\beta} \,|\, \boldsymbol{\alpha}, \mathbf{D}_\beta, \mathbf{h}^2, h_e^2, \sigma^2$, integrating out $\mathbf{u}$.
4. Sample $\mathbf{D}_\beta, \mathbf{h}^2, h_e^2 \,|\, \boldsymbol{\alpha}, \boldsymbol{\beta}, \sigma^2$, integrating out $\mathbf{u}$.
5. Sample $\mathbf{u} \,|\, \boldsymbol{\alpha}, \boldsymbol{\beta}, \mathbf{h}^2, h_e^2, \sigma^2$.

By caching intermediate calculations, the most expensive calculation for all five steps is a single Cholesky decomposition of a square matrix with $\min(b, n)$ rows.

We describe each step in detail below. First, we define several quantities that are re-used in multiple steps: $\mathbf{V}(\mathbf{h}^2) = \mathbf{Z}\mathbf{K}(\mathbf{h}^2)\mathbf{Z}^\intercal + h_e^2\mathbf{I}$ is the covariance of $\mathbf{y}$ from the random effects as a function of $\mathbf{h}^2$; and $\mathbf{V}_\beta = \mathbf{X}_2\mathbf{D}_\beta\mathbf{X}_2^\intercal + \mathbf{V}(\mathbf{h}^2)$ is the covariance of $\mathbf{y}$ after integrating out the regularized regression coefficients $\boldsymbol{\beta}$. Sampling steps in Gibbs sampler are as follows:

1. We assume that $a << n$. The conditional posterior of $\boldsymbol{\alpha}$ is given as

$$\begin{aligned} \boldsymbol{\alpha} \,|\, \cdot &\sim \mathcal{N}(\mathbf{A}_\alpha^{-1}\mathbf{X}_1^\intercal\mathbf{V}_\beta^{-1}\mathbf{y}, \sigma^2\mathbf{A}_\alpha^{-1}) \\ \mathbf{A}_\alpha &= \mathbf{X}_1^\intercal\mathbf{V}_\beta^{-1}\mathbf{X}_1 \end{aligned} \tag{S6}$$

   Inverting the $n \times n$ matrix $\mathbf{V}_\beta$ is expensive. We describe tricks to speed up this calculation below. Note that the $\mathbf{A}_\alpha$ is a small $a \times a$ matrix, so its Cholesky decomposition is inexpensive.

2. This step follows Makalic and Schmidt (2016) and modified for correlated residuals and un-regularized coefficients $\boldsymbol{\alpha}$. The conditional posterior of $\sigma^2$ is given as

$$\sigma^2 \,|\, \cdot \sim \mathrm{iG}(a_0 + n/2, b_0 + \boldsymbol{\varepsilon}^\intercal\mathbf{V}_\beta^{-1}\boldsymbol{\varepsilon}/2) \tag{S7}$$

   where $\boldsymbol{\varepsilon} = \mathbf{y} - \mathbf{X}_1\boldsymbol{\alpha}$ and $\mathbf{V}_\beta^{-1}$ is re-used from the previous step above.

3. This step also follows Makalic and Schmidt (2016) where

$$\begin{aligned} \boldsymbol{\beta}|\cdot &\sim \mathcal{N}(\mathbf{A}_\beta^{-1}\mathbf{X}_2^\intercal\mathbf{V}(\mathbf{h}^2)^{-1}\boldsymbol{\varepsilon}, \sigma^2\mathbf{A}_\beta^{-1}) \\ \mathbf{A}_\beta &= \mathbf{X}_2^\intercal\mathbf{V}(\mathbf{h}^2)^{-1}\mathbf{X}_2 + \mathbf{D}_\beta^{-1} \end{aligned} \tag{S8}$$

   where $\boldsymbol{\varepsilon}$ is defined as above. This step requires calculating $\mathbf{V}(\mathbf{h}^2)^{-1}$ and $\mathbf{A}_\beta^{-1}$. The former is $n \times n$ matrix, and the latter is a $b \times b$ matrix. We discuss below how these calculations can be accelerated, particularly when $b > n$.

4. Updates for $\mathbf{D}_\beta$ and $\mathbf{h}^2$ are independent. Once $\mathbf{h}^2$ is updated, $h_e^2$ is calculated as $1 - \sum_i h_i^2$.

   (a) We use the horseshoe prior form as a default for $\mathbf{D}_\beta = \mathrm{diag}(\phi_i^2\tau^2)$, specified as:

$$\begin{aligned} \beta_i \,|\, \sigma^2 &\sim \mathrm{N}(0, \phi_i^2\tau^2\sigma^2) \\ \phi_i &\sim \mathcal{C}^+(0,1) \\ \tau &\sim \mathcal{C}^+(0,\tau_0). \end{aligned} \tag{S9}$$

   We sample the parameters $\phi_i$ and $\tau$ using the algorithm of Makalic and Schmidt (2016) by introducing two new variables $\nu_i$ and $\xi$ such that

$$\begin{aligned} \phi_i^2 \,|\, \nu_i &\sim \mathrm{iG}(1/2, 1/\nu_i), \qquad \nu_i \sim \mathrm{iG}(1/2, 1) \\ \tau^2 \,|\, \xi &\sim \mathrm{iG}(1/2, 1/\xi), \qquad \xi \sim \mathrm{iG}(1/2, 1/\tau_0^2). \end{aligned} \tag{S10}$$

   Now, we can sample these parameters using the following steps:

   i. $\phi_i^2 \,|\, \cdot \sim \mathrm{iG}(1, \nu_i^{-1} + \beta_i^2/(2\tau^2\sigma^2))$
   ii. $\nu_i \,|\, \cdot \sim \mathrm{iG}(1, 1 + 1/\phi_i^2)$
   iii. $\tau^2 \,|\, \cdot \sim \mathrm{iG}\Big((b+1)/2, \xi^{-1} + (2\sigma^2)^{-1}\sum_{i=1}^b \beta_i^2\phi_i^{-2}\Big)$
   iv. $\xi \,|\, \cdot \sim \mathrm{iG}(1, 1/\tau^2 + 1/\tau_0^2)$

However, recall that when sampling using columns of $\mathbf{Y}$, $\boldsymbol{\beta}$ contains a column of $\boldsymbol{\Lambda}$ and a column of $\mathbf{B}_{2R}$ (i.e. $\boldsymbol{\beta} = [\boldsymbol{\lambda}_j^\intercal, \mathbf{b}_{2Rj}^T]^\intercal$). When sampling using columns of $\mathbf{F}$, $\boldsymbol{\beta}$ only contains a column of $\mathbf{B}_{2F}$ (i.e. $\boldsymbol{\beta} = \mathbf{b}_{2Fk}$). The global shrinkage parameter $\tau$ differs between elements of $\boldsymbol{\Lambda}$ and elements of $\mathbf{B}_{2R}$ and $\mathbf{B}_{2F}$ because global shrinkage applies to the whole matrix $\boldsymbol{\Lambda}$, but is unique to individual columns of $\mathbf{B}_{2R}$ and $\mathbf{B}_{2F}$. Also, the elements of $\boldsymbol{\Lambda}$ have the additional factor $\delta_k$ in their prior variance which increases the shrinkage from row $k$ to row $k+1$ (equation S2).

Therefore, we sample a different $\tau$ for all elements of $\boldsymbol{\Lambda}$ which is independent of the $\tau_{b_{2Rj}}$ and $\tau_{b_{2Fk}}$. Sampling this parameter requires all elements of $\boldsymbol{\Lambda}$ (not just those corresponding to the current column of $\mathbf{Y}$). The update for $\tau$ is:

$$\tau^2 \,|\, \cdot \sim \text{iG}\left(\frac{Kt+1}{2}, \frac{1}{\xi} + \frac{1}{2}\sum_{k=1}^{K}\left[\prod_{l=1}^{k}\frac{1}{\delta_l}\right]\sum_{j=1}^{t}\frac{\lambda_{kj}^2}{\phi_{kj}^2\sigma_{Rj}^2}\right). \tag{S11}$$

Finally, the update for $\delta_h$ is:

$$\delta_h \,|\, \cdot \sim \text{iG}\left(a_\delta + \frac{t}{2}(K-h+1), b_\delta + \frac{1}{2\tau^2}\sum_{k=h}^{K}\left[\prod_{l=1, l\neq h}^{k}\frac{1}{\delta_l}\right]\sum_{j=1}^{t}\frac{\lambda_{kj}^2}{\phi_{kj}^2\sigma_{Rj}^2}\right) \tag{S12}$$

which also depends on the whole matrix $\boldsymbol{\Lambda}$. The Gibbs updates for $\tau$ and $\delta_h$ tend to mix poorly because of their strong dependence. Since these updates are relatively inexpensive, we repeat these two steps 100 times in a row per overall iteration of the MCMC chain to improve mixing.

(b) The prior for $\mathbf{h}^2$ is discrete on the $M$-dimensional simplex (for $M$ random effects). We use a Metropolis-Hastings step to update $\mathbf{h}^2$. We propose a new value $\mathbf{h}^2_{(1)}$ uniformly from the set of all values with $||\mathbf{h}^2_{(1)} - \mathbf{h}^2_{(0)}||_2 < \epsilon$, and then calculate:

$$r = \frac{p(\mathbf{h}^2_{(1)} \,|\, \cdot)g(\mathbf{h}^2_{(0)} \,|\, \mathbf{h}^2_{(1)})}{p(\mathbf{h}^2_{(0)} \,|\, \cdot)g(\mathbf{h}^2_{(1)} \,|\, \mathbf{h}^2_{(0)})} \tag{S13}$$

where the transition probability $g(\mathbf{h}^2_{(i)} \,|\, \mathbf{h}^2_{(j)})$ is proportional to the number of grid cells within $\epsilon$ of $\mathbf{h}^2_{(j)}$, and

$$p(\mathbf{h}^2_{(i)} \,|\, \cdot) \propto \left|\mathbf{V}(\mathbf{h}^2_{(i)})\right|^{-1/2} \times \exp\left\{-\frac{1}{2\sigma^2}\boldsymbol{\varepsilon}^{*\intercal}\mathbf{V}(\mathbf{h}^2_{(i)})^{-1}\boldsymbol{\varepsilon}^*\right\} \times p(\mathbf{h}^2_{(i)})$$

with $\boldsymbol{\varepsilon}^* = \mathbf{y} - \mathbf{X}_1\boldsymbol{\alpha} - \mathbf{X}_2\boldsymbol{\beta}$. We then accept $\mathbf{h}^2_{(1)}$ with probability $\min(1, r)$. The most expensive part of this calculation is evaluating the determinant and inverse of the $n \times n$ matrix $\mathbf{V}(\mathbf{h}^2_{(i)})$, which we solve through a Cholesky decomposition. Since $\mathbf{h}^2$ can take only a discrete set of values, we simply pre-calculate and cache all possible decompositions before starting the MCMC and re-use them throughout the chain for all $j = 1, \ldots, p$ $\mathbf{y}$ vectors.

(c) The update for $\mathbf{u}$ is given as the following

$$\begin{aligned} \mathbf{u} \,|\, \cdot &\sim \mathcal{N}\big(\mathbf{A}_\mathbf{u}(\mathbf{h}^2)^{-1}\mathbf{Z}^\intercal\boldsymbol{\varepsilon}^*/h_e^2\sigma^2, \mathbf{A}_\mathbf{u}(\mathbf{h}^2)^{-1}\big) \\ \mathbf{A}_\mathbf{u}(\mathbf{h}^2) &= \frac{1}{\sigma^2}\left(\frac{1}{h_e^2}\mathbf{Z}^\intercal\mathbf{Z} + \mathbf{V}(\mathbf{h}^2)^{-1}\right) \end{aligned} \tag{S14}$$

Here, $\mathbf{A_u}(\mathbf{h}^2)$ is an $r \times r$ matrix (where $r = \sum_{m=1}^{M} r_m$) which is a function of $\mathbf{h}^2$ and can be much larger than $n \times n$ if there are multiple full-rank random effects. Again, rather than calculating the Cholesky decomposition of $\mathbf{A_u}$ during each iteration for each trait, we note that there are a finite number of these matrices (indexed by $\mathbf{h}^2$) and pre-calculate and cache all before running the MCMC. Generally these matrices are sparse and can be stored efficiently.

#### 1.4 Opportunities for Caching Expensive Calculations

We noted above that we can cache Cholesky decompositions for each $\mathbf{V}(\mathbf{h}^2)$ and $\mathbf{A_u}(\mathbf{h}^2)$ indexed by $\mathbf{h}^2$ and re-use them for all $p$ traits and all iterations of the MCMC chain. Additionally, in Steps 1-3, the matrices $\mathbf{V}_\beta^{-1}$ and $\mathbf{A}_\beta^{-1}$ are closely related through the Binomial Inverse Theorem where

$$\begin{aligned}\mathbf{V}_\beta^{-1} &= [\mathbf{X}_2\mathbf{D}_\beta\mathbf{X}_2^\intercal + \mathbf{V}(\mathbf{h}^2)]^{-1} \\ &= \mathbf{V}(\mathbf{h}^2)^{-1} - \mathbf{V}(\mathbf{h}^2)^{-1}\mathbf{X}_2\left[\mathbf{X}_2^\intercal\mathbf{V}(\mathbf{h}^2)^{-1}\mathbf{X}_2 + \mathbf{D}_\beta^{-1}\right]^{-1}\mathbf{X}_2^\intercal\mathbf{V}(\mathbf{h}^2)^{-1} \\ &= \mathbf{V}(\mathbf{h}^2)^{-1} - \mathbf{V}(\mathbf{h}^2)^{-1}\mathbf{X}_2\mathbf{A}_\beta^{-1}\mathbf{X}_2^\intercal\mathbf{V}(\mathbf{h}^2)^{-1}/\sigma^2.\end{aligned} \tag{S15}$$

Since $\mathbf{V}(\mathbf{h}^2)^{-1}$ is pre-calculated, we only need to calculate the smaller of the $\mathbf{V}_\beta^{-1}$ and $\mathbf{A}_\beta^{-1}$ matrices and then form the other through matrix multiplications. Because we need the Cholesky decomposition $\mathbf{L}_\beta\mathbf{L}_\beta^\intercal = \mathbf{A}_\beta$ to sample $\boldsymbol{\beta}$ in Step 3, if $b < n$, we can calculate this directly and then use $\mathbf{L}_\beta$ to calculate $\mathbf{V}_\beta^{-1}$ for Steps 1 and 2. However, if $b > n$, we instead sample $\boldsymbol{\beta}$ using a modified version of the algorithm demonstrated by Bhattacharya *et al.* (2016). Let $\mathbf{L}\mathbf{L}^\intercal$ be the Cholesky decomposition of $\mathbf{V}(\mathbf{h}^2)$. Then

1. Sample $\mathbf{a} \sim \mathcal{N}(\mathbf{0}, \sigma^2\mathbf{D}_\beta)$ and $\mathbf{d} \sim \mathcal{N}(\mathbf{0}, \mathbf{I}_n)$ independently.
2. Set $\mathbf{v} = 1/\sigma\mathbf{L}^{-1}\mathbf{X}_2\mathbf{a} + \mathbf{d}$
3. Set $\mathbf{A}_w = \mathbf{L}^{-1}\mathbf{X}_2\mathbf{D}_\beta\mathbf{X}_2^\intercal\mathbf{L}^{-\intercal} + \mathbf{I}_n$
4. Solve $\mathbf{A}_w\mathbf{w} = (\mathbf{L}^{-1}\tilde{\mathbf{y}}/\sigma - \mathbf{v})$ to obtain $\mathbf{w}$.
5. Set $\boldsymbol{\beta} = \mathbf{a} + \sigma\mathbf{D}_\beta\mathbf{X}_2^\intercal\mathbf{L}^{-\intercal}\mathbf{w}$.

Note that $\mathbf{L}\mathbf{A}_w\mathbf{L}^\intercal = \mathbf{V}_\beta$. If we have already calculated $\mathbf{V}_\beta^{-1}$, we can simplify these steps to:

1. Sample $\mathbf{a} \sim \mathcal{N}(\mathbf{0}, \sigma^2\mathbf{D}_\beta)$ and $\mathbf{d} \sim \mathcal{N}(\mathbf{0}, \mathbf{I}_n)$ independently.
2. Set $\mathbf{v}^* = 1/\sigma\mathbf{X}_2\mathbf{a} + \mathbf{L}\mathbf{d}$
3. Solve $\mathbf{V}_\beta\mathbf{w}^* = (\tilde{\mathbf{y}}/\sigma - \mathbf{v}^*)$ to obtain $\mathbf{w}^*$.
4. Set $\boldsymbol{\beta} = \mathbf{a} + \sigma\mathbf{D}_\beta\mathbf{X}_2^\intercal\mathbf{w}^*$.

Finally, if $\mathbf{X}_2$ has row-rank less than $n$, we can factor as $\mathbf{U}\mathbf{V}^\intercal$ where $\mathbf{U}$ and $\mathbf{V}$ are $n \times m$ and $m \times b$ matrices, respectively, with $m < n$. This often occurs if $m$ genotypes are repeated multiple times in the same dataset. In this case, we can speed up the calculation of $\mathbf{V}_\beta^{-1}$ using the Binomial Inverse Theorem:

$$\begin{aligned}\mathbf{V}_\beta^{-1} &= \left[\mathbf{X}_2\mathbf{D}_\beta\mathbf{X}_2^\intercal + \mathbf{V}(\mathbf{h}^2)\right]^{-1} \\ &= \left[\mathbf{U}\mathbf{V}\mathbf{D}_\beta\mathbf{V}^\intercal\mathbf{U}^\intercal + \mathbf{V}(\mathbf{h}^2)\right]^{-1} \\ &= \mathbf{V}(\mathbf{h}^2)^{-1} - \mathbf{V}(\mathbf{h}^2)^{-1}\mathbf{U}\left[(\mathbf{V}\mathbf{D}_\beta\mathbf{V}^\intercal)^{-1} + \mathbf{U}^\intercal\mathbf{V}(\mathbf{h}^2)^{-1}\mathbf{U}\right]^{-1}\mathbf{U}^\intercal\mathbf{V}(\mathbf{h}^2)^{-1}\end{aligned} \tag{S16}$$

This involves only two $m \times m$ matrix inversions, which is beneficial if $m << n$. Further caching is possible to prevent redundant matrix-matrix multiplications. Since $\mathbf{X}_2$ is constant across the $p$ traits, the terms $\mathbf{L}^{-1}\mathbf{X}_2$ and $\mathbf{X}_2^\intercal\mathbf{V}(\mathbf{h}^2)^{-1}\mathbf{X}_2$, (or $\mathbf{L}^{-1}\mathbf{U}$ and $\mathbf{U}^\intercal\mathbf{V}(\mathbf{h}^2)^{-1}\mathbf{U}$) are also constant for any two traits that share the same value for $\mathbf{h}^2$. This can dramatically reduce the computational complexity when $p$ is large.

#### 1.5 Blocking missing data

Most existing Gibbs samplers for factor models require complete observations because they condition on the whole trait vectors for each individual and, therefore, must impute any missing data. While data imputation is straightforward in Bayesian models (missing data is treated as an unknown parameter and included in the MCMC), conditioning on imputed values in a Gibbs sampler leads to very long-term autocorrelations in MCMC chains. `MegaLMM` largely avoids this by dropping unobserved values from the likelihood. However, there is a trade-off between simplifying the likelihood and computational efficiency. In particular, the pre-cached Choleksy decompositions of the random effect covariance matrices and other intermediate calculations can only be applied to sets of traits with complete observations across the same individuals. If every trait had a unique pattern of missing observations, pre-caching would be extremely memory-intensive and inefficient.

In many data sets, distinct subsets of traits share approximately the same set of missing observations. For example, all agronomic traits may be measured on a subset of lines in a breeding trial, while hyperspectral reflectance is measured on all lines. Or a similar subset of all possible lines may be grown in nearby fields of a multi-environment trial.

In these cases, we partition the full matrix of traits $\mathbf{Y}$ into a list of smaller trait matrices $\{\widetilde{\mathbf{Y}}_1, \ldots, \widetilde{\mathbf{Y}}_S\}$, where each sub-matrix $\widetilde{\mathbf{Y}}_s$, contains only those individuals with observations for this subset of the traits. We select the partitions by attempting to minimizing the number of unobserved values within each $\widetilde{\mathbf{Y}}_s$, for a given number of partitions, using a sequence of binary partitions of the original trait matrix. We impute values for all missing data in $\mathbf{Y}$ during Step 4, but only condition on the imputed values in each $\widetilde{\mathbf{Y}}_s$ during Steps 1 and 3. This greatly reduces autocorrelation in the MCMC while minimizing the number of pre-cached intermediate calculations that need to be stored.

#### 1.6 Further Mixing Issues

As in any factor model, the factor loadings in our model are not identifiable in the likelihood. However, the horseshoe prior on elements of $\boldsymbol{\Lambda}$ does help make these parameters identifiable in the posterior (except for sign flips). In general, we find that coefficients of each row $\boldsymbol{\lambda}_k$ mix reasonably well. It is important to note that the ordering of the rows in $\boldsymbol{\Lambda}$ from most-to-least important does not mix effectively. While the correct ordering is important to ensure that important factors are not shrunk too much, we are generally not interested in the order *per se* as much as the identities of each factor, and we find that genomic prediction outcomes are relatively robust to mixing issues of factor order. To improve model convergence, during the burn-in period, we periodically stop the MCMC chain, assess the order of the factors, and manually re-order the factors based on current estimates of $m_k \,|\, \tau_k$. We also prune factors when when the pairwise correlations among columns of $\mathbf{F}$ are too high ($\rho > 0.6$).

#### 1.7 Additional strategies for computational efficiency.

When there is only one random effect in our model, we can gain further computational efficiency by diagonalizing the model where we pre-multiply both sides of Eq. (2) by the transpose of the eigenvectors from $\mathbf{K}$. This makes $\mathbf{V}(\mathbf{h}^2)$ diagonal for all values of $\mathbf{h}^2$. By storing the Cholesky decomposition of these diagonal matrices as sparse objects, we can take advantage of efficient sparse linear algebra libraries for dramatic gains in efficiency. This strategy only works when $\mathbf{Y}$ is without missing data. However, we can modify the strategy slightly by calculating eigenvalue decompositions of $\widetilde{\mathbf{K}}_s$ for each subset of the partitioned trait sub-matrices $\widetilde{\mathbf{Y}}_s$, and then pre-multiply each sub-matrix by the corresponding set of eigenvectors.

When more than one random effect is included, complete diagonalization is no longer possible and so we resort to pre-caching Cholesky decompositions of $\mathbf{V}(\mathbf{h}^2)$ for each value of $\mathbf{h}^2$. When random effects are low-rank, Cholesky decompositions of their covariance matrices can sometimes be stored as sparse matrices to reduce memory and computational demands. We check the number of zero-elements in each Cholesky decomposition to determine whether to store it as a sparse or dense matrix.

Finally, we code nearly all expensive linear algebra calculations in our `R` package in C++ using the `Eigen` library with `RcppEigen` (Bates and Eddelbuettel 2013), and parallelize calculations across traits whenever possible using OpenMP.

#### 2 Supplementary Figures

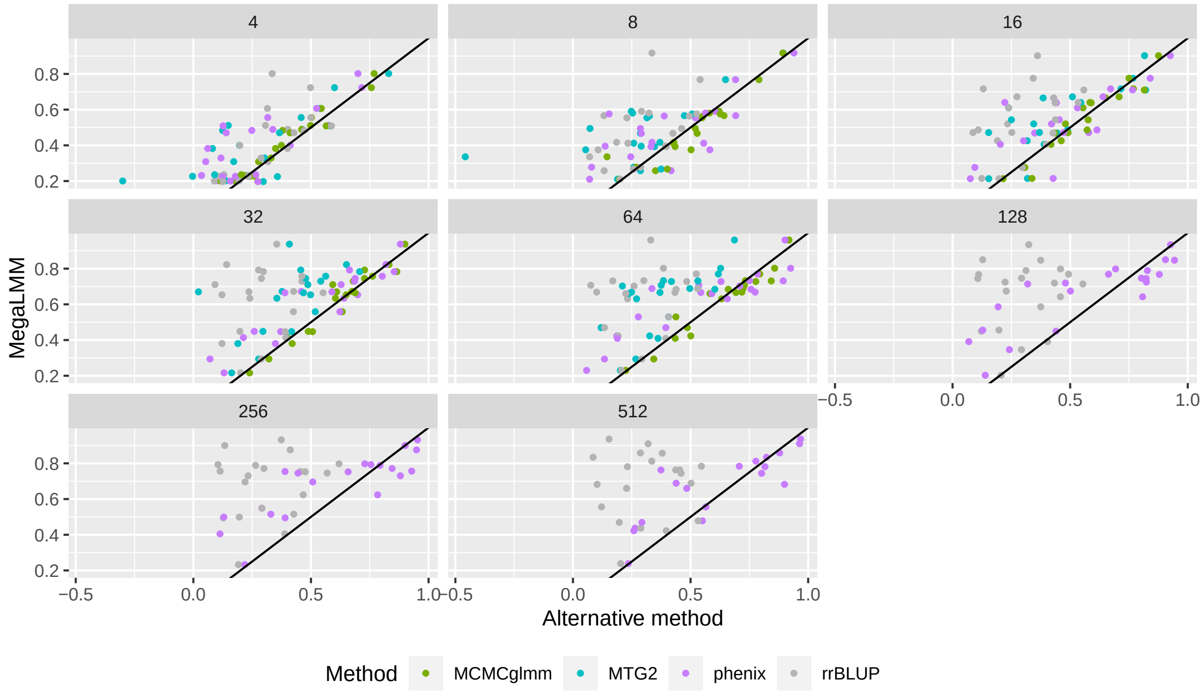

**Figure S1. Prediction accuracies for each target gene expression trait relative to `MegaLMM`.** Each panel shows the estimate accuracies ($\hat{\rho}_g(\hat{\mathbf{g}}, \mathbf{g})$ for each of the 20 target genes. The accuracy for `MegaLMM` is on the x-axis and the accuracy of the other 4 methods is on the y-axis. The black line shows equal accuracy. Points above the line show datasets where the accuracy of `MegaLMM` was greater than the alternative method. These are the same results as presented in Figure 2A.

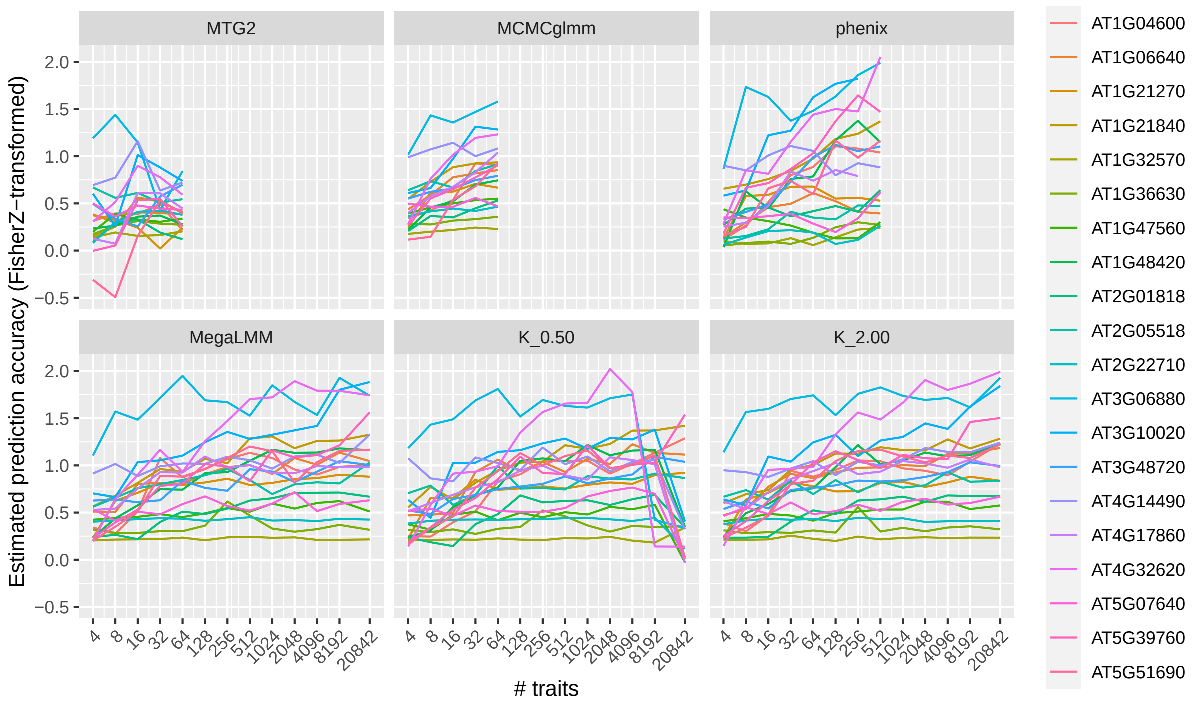

**Figure S2. Prediction accuracies for each target gene expression trait as a function of dataset size.** Each line traces the change in estimate prediction accuracy ($\hat{\rho}_g(\hat{\mathbf{g}}, \mathbf{g})$ for one of the 20 focal genes as more secondary traits are added to the dataset. Each panel shows a different method. Methods `K_0.50` and `K_2.00` are MegaLMM with $K$ set to either 0.5x or 2x the value used in Figure 2. These are the same results as presented in Figure 2A.

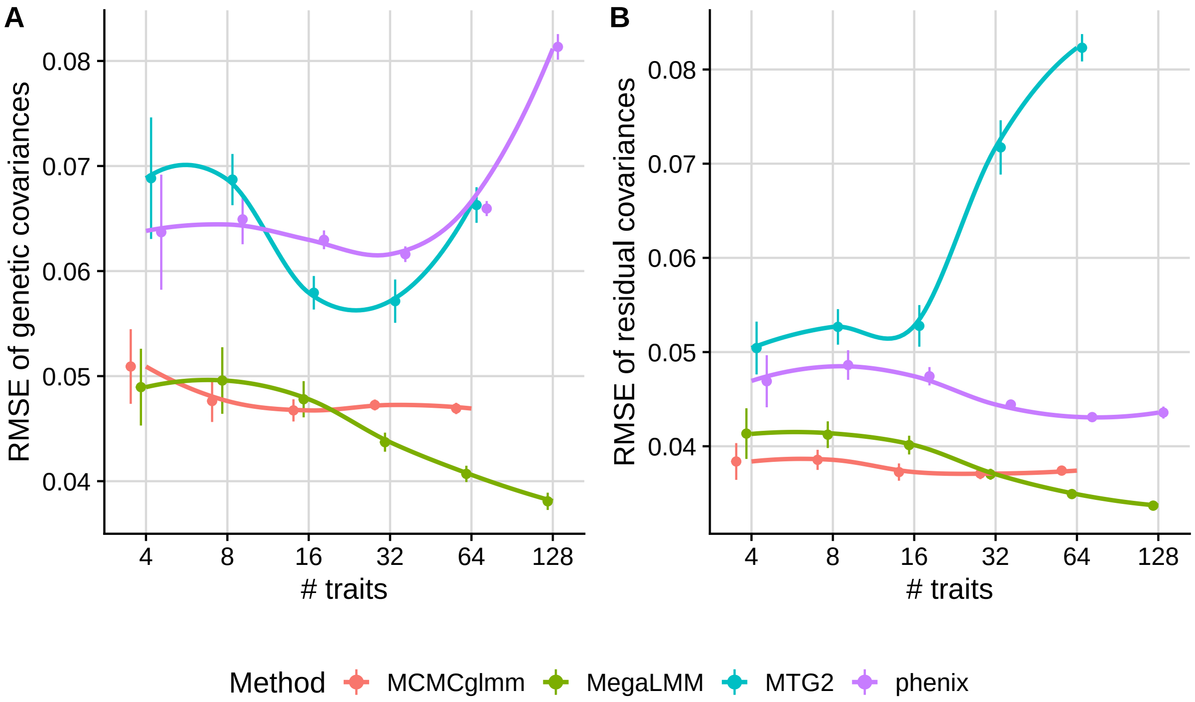

**Figure S3. Accuracy of covariance estimates in simulated data.** We generated 20 simulated datasets each with 128 genes based on the Arabidopsis gene expression data. In each dataset, we sampled two sets of 128 genes and used each to calculated an empirical correlation matrix. We converted the correlation matrices into genetic ($\mathbf{G}$) and residual ($\mathbf{R}$) covariance matrices by sampling a $h^2$ value independently from $[0.1, 0.8]$ for each gene, and then used the matrices and the genomic relationship matrix $\mathbf{K}$ to generate genetic and residual values for 128 traits. Using subsets of these simulated data with varying numbers of genes, we estimated genetic and residual covariances using each of the four comparison methods using the same parameters as in Figure 2, and then assessed the accuracy of covariance estimation by calculating the square-root of the mean squared error (RMSE) between each covariance value (excluding variances). The whole procedure was repeated 20 times with different simulated datasets. Points and bars show the mean and 1SE of the RMSE values for each method. Curves were estimated with 'scales::loess'.

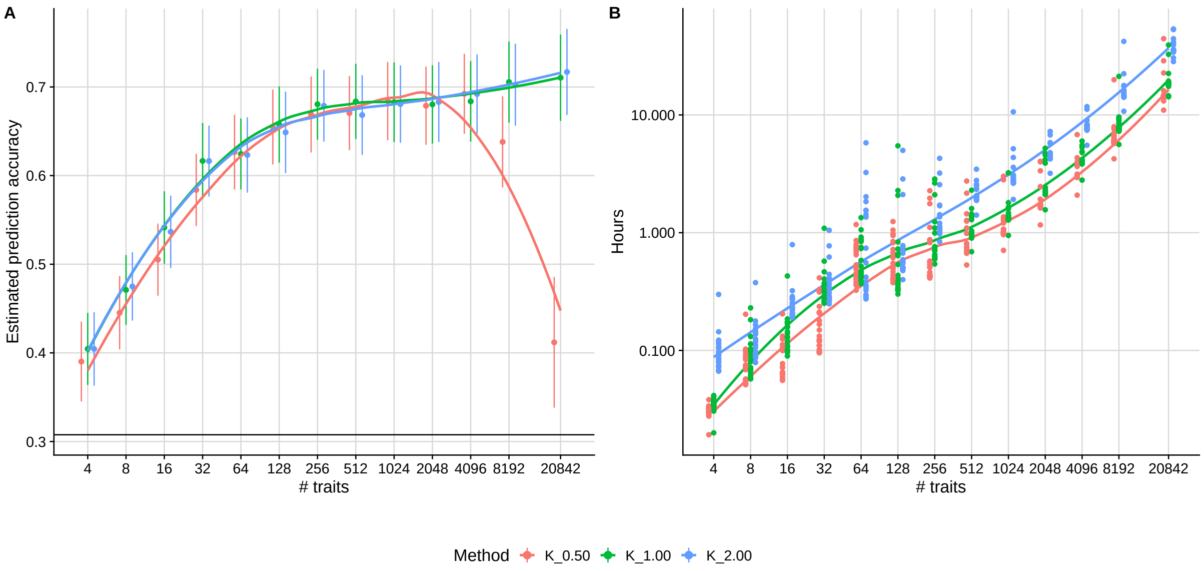

**Figure S4. Sensitivity of `MegaLMM` to changes in $K$.** This figure parallels Figure 2 from the main text except estimated prediction accuracies (**A**) and computational times (**B**) are shown for three runs of `MegaLMM` with varying $K$. For each dataset size, $K$ was either increased by a factor of 2 (`K_2.00`) or decreased by a factor of 2 (`K_0.50`) . Otherwise datasets and parameters are the same as for Figure 2.

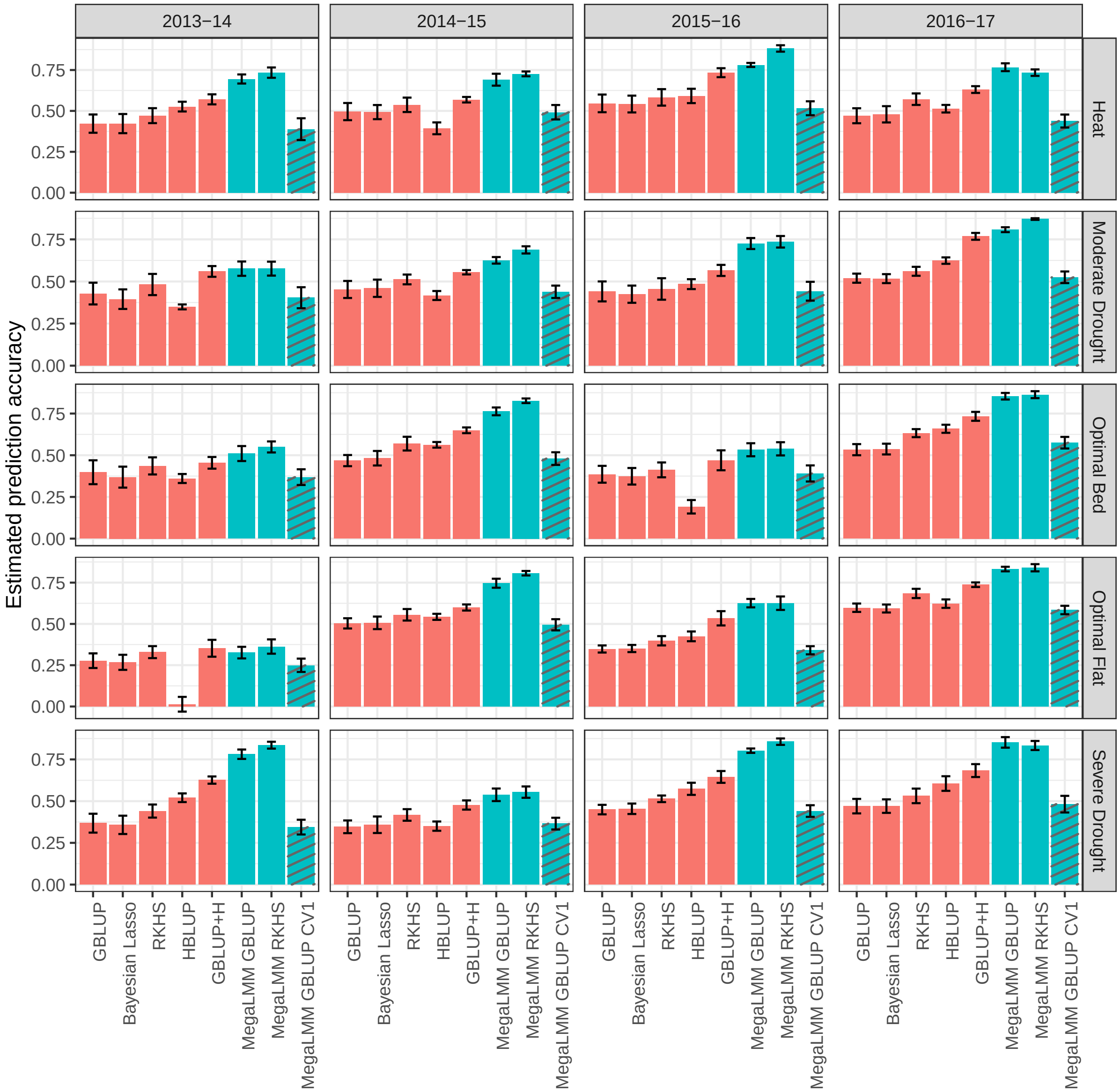

**Figure S5. Performance of single-trait and multi-trait genomic prediction for wheat yield across 20 experiments.** The same analysis reported in Figure 3 was repeated for the 19 other treatment:year datasets reported by Krause *et al.* (2019). In each panel, bars show the mean ± SE over five cross-validation runs, each with a unique 50:50 testing:training partition. The same partitions were used for each method in each run. `MegaLMM` methods are colored in blue. The hashed lines indicate a multivariate prediction using the CV1 approach (i.e. no secondary traits included in the testing data for prediction). The dataset in Figure 3 was 2014-2015 Optimal Bed.

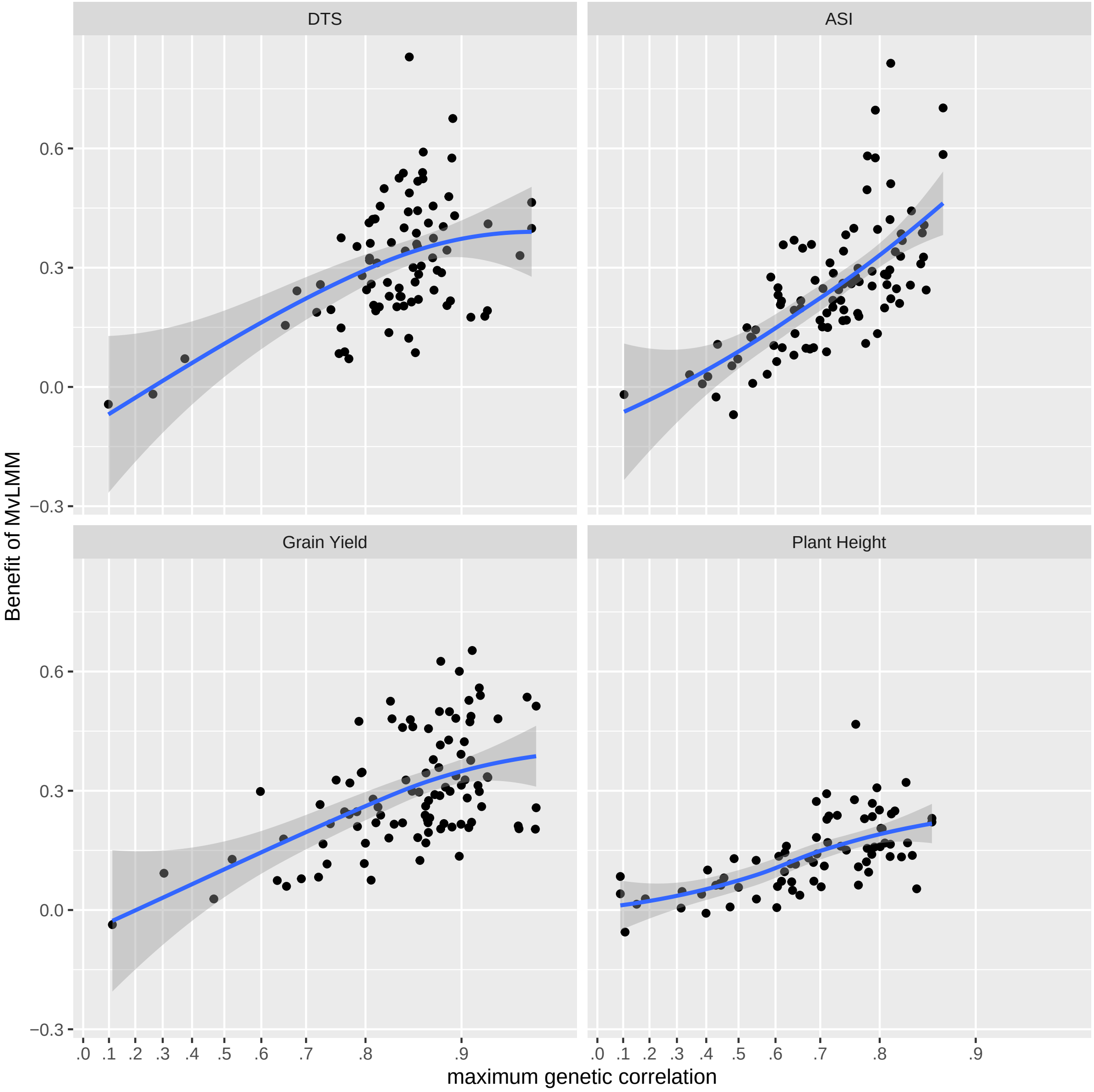

**Figure S6. Relationship between genetic correlation and benefit of MvLMM across Genomes2Fields site-years.** In each panel, each point represents a single site-year. The x-axis shows the maximum genetic correlation between the trait values for that site-year and all other site-years. The y-axis is the difference in the Fisher Z-transformed estimates of genomic prediction accuracy between the full MvLMM (implemented in `MegaLMM`) and a univariate GBLUP model (implemented in `rrBLUP`). Traits included: days to silking (DTS), anthesis-silking interval (ASI), grain yield, and plant height.

### 3 Supplementary Tables

**Table S1. Default hyperparameters for user-customizable prior distributions.** Default values provided in the `MegaLMM` R package are provided. These values were used for the reported analyses unless otherwise noted.

| Parameter | Distribution | Hyperparameters | Interpretation |
|---|---|---|---|
| $\sigma^2_{Rj}$ | $\mathrm{iG}(\alpha = \nu - 1, \beta = 1/V\nu)$ | $\nu = 10, V = 0.5$ | The variance of each column of $\mathbf{E}_R$ has mean $\approx V$, with spread determined by $\nu$. |
| $\tau$ | $\mathcal{C}^+(0, \pi_0/n(1-\pi_0)^2)$ | $\pi_0 = 0.1$ | The proportion of effectively non-zero elements in the first row of $\mathbf{\Lambda}$ is $\pi_0$. |
| $\delta_h$ | $iG(a_\delta, b_\delta)$ | $a_\delta = 3, b_\delta = 1$ | The expected odds of being non-zero for an element of $(\mathbf{\Lambda}^\intercal)_{h+1}$ decreases by a factor of $\approx a_\delta/b_\delta$ relative to $(\mathbf{\Lambda}^\intercal)_h$. |
| $\tau_{\mathbf{b}_{2R}}, \tau_{\mathbf{b}_{2F}}$ | $\mathcal{C}^+(0, \pi_0/n(1-\pi_0)^2)$ | $\pi_0 = 0.1$ | The proportion of effectively non-zero elements in the first row of $\mathbf{B}_{2R}$ or $\mathbf{B}_{2F}$ is $\pi_0$. |
| $\mathbf{h}^2_{Rj}, \mathbf{h}^2_{Fj}$ | $1/l$ | $l = 20$ | Uniform over $l$ equally spaced grid cells on the unit simplex. When $M > 1$, $l$ counts the number of valid grid cells so is less than $l^M$. |
